# Direct Relationship between Protein Expression and Progeny Yield of Herpes Simplex Virus 1 Unveils a Rate-limiting Step for Virus Production

**DOI:** 10.1101/2023.06.07.544155

**Authors:** Moeka Nobe, Yuhei Maruzuru, Kosuke Takeshima, Fumio Maeda, Hideo Kusano, Raiki Yoshimura, Takara Nishiyama, Hyeongki Park, Yoshitaka Kozaki, Shingo Iwami, Naoto Koyanagi, Akihisa Kato, Tohru Natsume, Shungo Adachi, Yasushi Kawaguchi

## Abstract

Although viral protein expression and progeny virus production were independently shown to be highly heterogenous in individual cells, their direct relationship, analyzed by considering their heterogeneities, has not been investigated to date. This study established a system to fractionate cells infected with a herpesvirus based on the levels of the global expression of viral late proteins, which are largely virion structural proteins, and to titrate virus yields in these fractions. This system demonstrated a direct relationship and indicated there was a threshold for the levels of viral late protein expression for progeny virus production and suggested that viral DNA cleavage/packaging was a rate-limiting step for progeny virus production. These findings, which were masked in previous studies performed at the entire population level, have uncovered a sophisticated viral strategy for efficient progeny virus production and shed new light on an effective target for the development of anti-viral drugs.

## INTRODUCTION

The state of viral gene expression has long been thought to be one of the critical determinants for virus progeny production ^1^. Viral infection is usually studied at the entire population level by averaging the outcomes of infection from each of large numbers of individual cells. In these previous studies, a relationship between viral gene expression and virus progeny production has necessarily been investigated based on classical time course experiments in which levels of viral gene expression and progeny virus titers were compared at various times after infection. Such studies demonstrated that the levels of viral gene expression, including the expressions of viral mRNA and protein, correlated well with progeny virus yields ^2–6^. However, accumulating evidence has indicated that viral infection at the single-cell or subpopulation level is highly heterogeneous, which has been masked in studies performed at the entire population level. Thus, it was reported that the progeny virus yield from individual cells spanned several orders of magnitude ^7–14^. Classical fluorescence microscopy and recent advances in single-cell RNA-sequencing have enabled the investigation of the state of viral gene expression at the single-cell level, revealing high heterogeneity in individual cells and identifying new subpopulations of infected cells with similar viral gene expression profiles ^15–17^. However, there is a lack of information on viral protein expression and virus progeny production in the same individual cell or subpopulation. Therefore, the direct relationship between viral protein expression and progeny virus production, analyzed by considering their heterogeneities at single cell or subpopulation levels, remains to be elucidated. These observations raised a fundamental question as to whether the well-established correlation between viral protein expression and progeny virus production detected at the entire population level by classical time-course experiments ^4,6^ does reflect a direct relationship between them.

Herpes simplex virus 1 (HSV-1), an extensively studied DNA virus, is a ubiquitous human pathogen, causing a variety of diseases including encephalitis, keratitis, and mucocutaneous and skin diseases including herpes labialis, genital herpes and herpetic whitlow ^18^. HSV-1 encodes more than 100 different proteins ^19–21^ and HSV-1 genes fall into three major classes: immediate-early (IE), early (E) and late (L), whose expressions are coordinately regulated and sequentially ordered in a cascade during lytic infection ^19^. Although there are some exceptions, virion structural proteins are largely encoded by L genes ^19^. Replication of the HSV-1 genome and packaging of the replicated viral genomes into nascent capsids occurs in the nucleus ^19^. Nascent nucleocapsids are exported to the cytoplasm through the perinuclear space between the inner nuclear membrane and outer nuclear membrane by a nuclear pore-independent and sequential envelopment/de-envelopment process ^19^. In the cytoplasm, capsids acquire a final envelope by budding into cytoplasmic vesicles and become infectious ^19^.

In this study, we constructed a reporter HSV-1 to monitor the global expression of viral L proteins by analyzing direct and quantitative relationship between L protein expression levels and virus progeny yields by sorting infected cells into subpopulations based on reporter gene expression levels and by determining virus titers in the subpopulations. The clarified relationship indicated a threshold in the levels of HSV-1 L protein expression for progeny virus production, suggesting viral DNA cleavage/packaging is a rate-limiting step for progeny virus production.

## RESULTS

### Construction and characterization of a reporter HSV-1 to analyze viral L protein expression and virus progeny yields at the subpopulation level

We constructed a recombinant virus rICP47/vUs11 expressing IE protein ICP47 and L protein Us11 fused to monomeric fluorescent proteins, TagRFP and VenusA206K (TagRFP-ICP47 and Venus-Us11), respectively (Fig. 1A and B). The growth of rICP47/vUs11 was similar to wild-type HSV-1(F) in HeLa cells at a multiplicity of infection (MOI) of 5 and reached a plateau 24 h after infection (Fig. 1C). Flow cytometric analyses showed 92% of HeLa cells inoculated with rICP47/vUs11 at an MOI of 5 were TagRFP positive 24 h after inoculation, indicating that in these experimental settings, most HeLa cells were infected with rICP47/vUs11 and HSV-1 gene expression was initiated in these infected cells (Fig. 1D and E). Frequencies of cells positive for Venus and TagRFP, or those positive for TagRFP and negative for Venus, were 67% and 25%, respectively (Fig. 1E). Cells positive for Venus and negative for TagRFP were barely detectable (Fig. 1D and E). Notably, the coefficient of variation (CV) for Venus-Us11 fluorescence intensities in each infected cell was significantly higher than for TagRFP-ICP47 fluorescence (Fig. 1F).

**Fig. 1.**
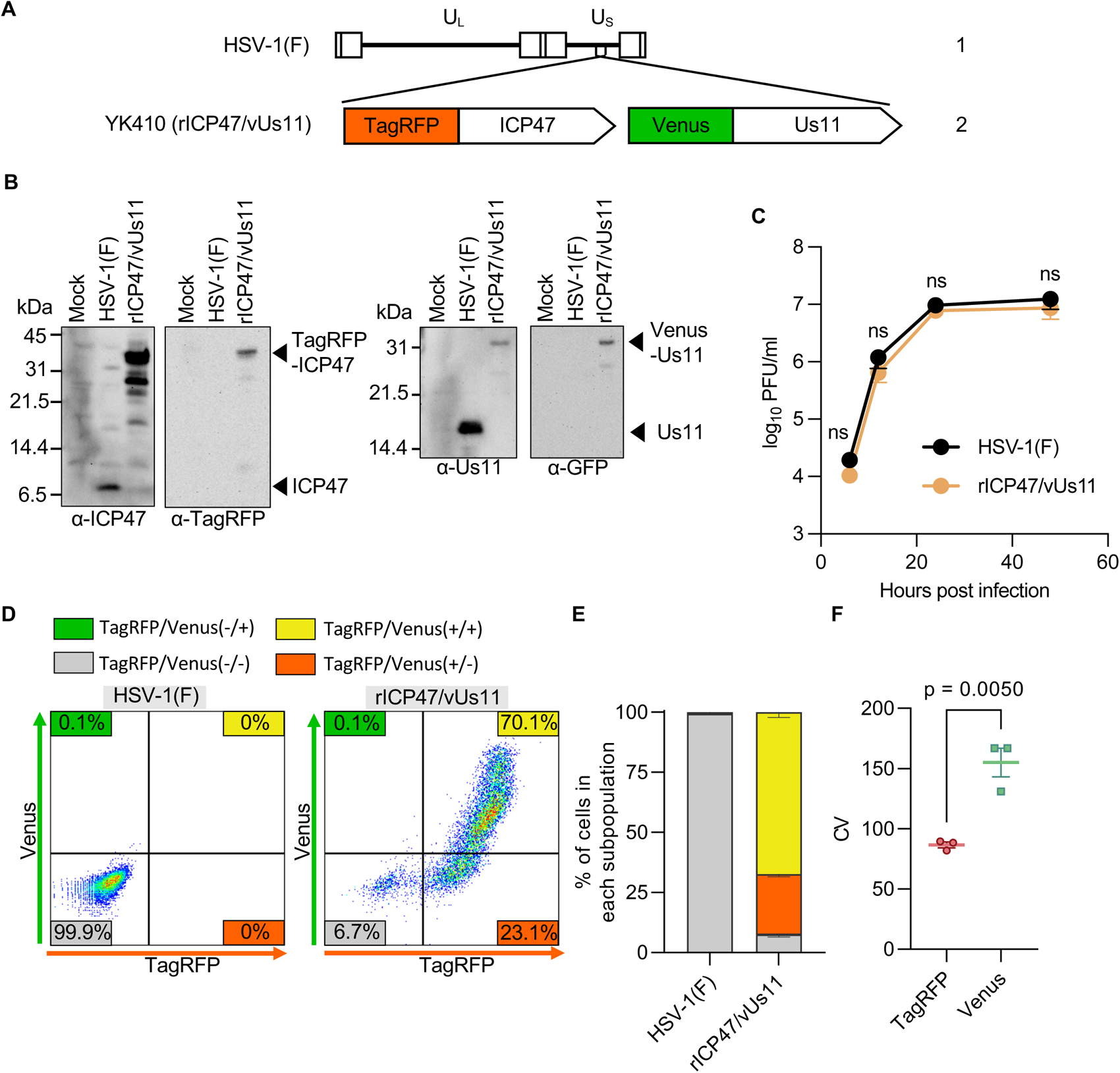
Characterization of the reporter HSV-1 rICP47/vUs11 generated in this study. (A) Schematic diagram of the genome structure of wild-type HSV-1(F) and rICP47/vUs11. Line 1, wild-type HSV-1(F) genome; line 2, domains of the Us11 and ICP47 coding regions. The positions of insertion of TagRFP and Venus are indicated. (B) HeLa cells mock-infected or infected for 24 h with wild-type HSV-1(F) or rICP47/vUs11 at an MOI of 5 were lysed and analyzed by immunoblotting with the indicated antibodies. (C) HeLa cells were infected at an MOI of 5 with wild-type HSV-1(F) or rICP47/vUs11. Each cell culture supernatant plus the infected cells were harvested at the indicated times post infection, and progeny viruses were assayed on Vero cells. (D) HeLa cells infected for 24 h with wild-type HSV-1(F) or rICP47/vUs11 at an MOI of 5 were analyzed by flow cytometry. (E) Quantitative bar graph of the proportion of cells in the TagRFP/Venus (+/+), TagRFP/Venus (+/-), TagRFP/Venus (-/+), and TagRFP/Venus (-/-) subpopulations shown in (D). (F) CV values for TagRFP-ICP47 and Venus-Us11 shown in (D). The data are representative of three independent experiments (B and D). Each value is the mean ± standard error of the results of four (C) or three (E and F) independent experiments. Statistical analysis was performed by unpaired Student’s *t*-test. ns, not significant (C and F).

To examine whether levels of Venus-Us11 fluorescence in rICP47/vUs11-infected cells were related to those of the global expression of HSV-1 L proteins, HeLa cells infected with rICP47/vUs11 at an MOI of 5 for 24 h were analyzed by flow cytometry and sorted into six subpopulations (f1 to f6) based on the levels of Venus-Us11 fluorescence intensities—Venus-Us11 fluorescence intensities increased in subpopulations from f1–f6 (S-Fig. 1A). The subpopulations were then subjected to LC-MS/MS to quantitate peptides of global HSV-1 L proteins (Fig. 2A and S-Fig. 1B). Among the HSV-1 L proteins detected (48 L proteins), there was a strong correlation between the levels of Venus-Us11 fluorescence intensity and the relative abundance of most HSV-1 L proteins including Venus-Us11 (33 L proteins, r>0.90, P<0.001; 10 L proteins, r>0.70, P<0.01; 4 L proteins, r>0.58, P<0.05) but not Us8.5 (r=0.54, P=0.07). These results suggested that Venus-Us11 fluorescence intensities reflected expression levels of global HSV-1 L proteins in rICP47/vUs11-infected cells.

**Fig. 2.**
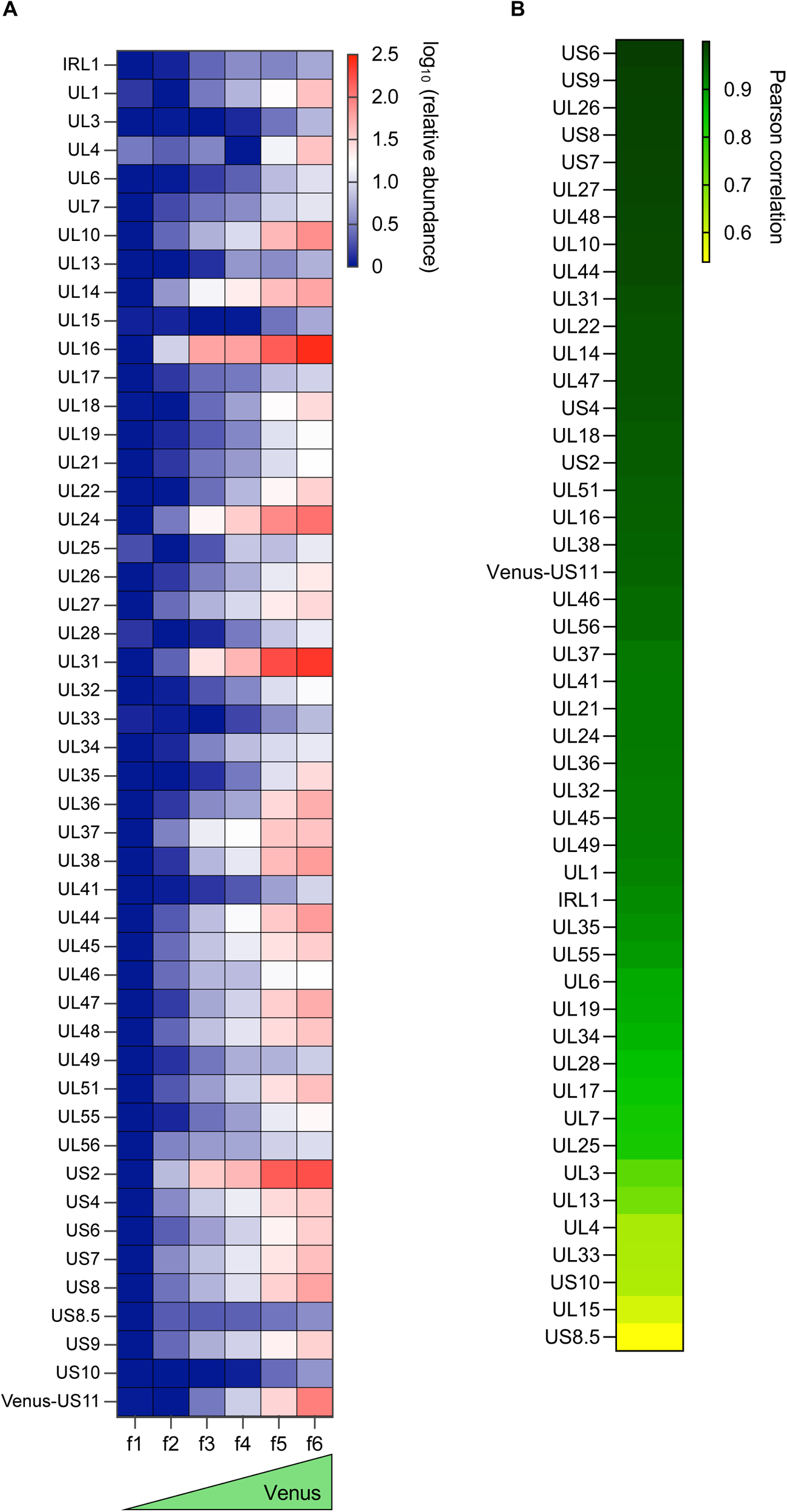
Abundance of HSV-1 L proteins in each subpopulation of rICP47/vUs11-infected cells. (A) HeLa cells were infected for 24 h with rICP47/vUs11 at an MOI of 5 and sorted into six subpopulations (f1 to f6) by cell sorting as shown in S-Fig. 1A and each subpopulation was analyzed by LC-MS/MS. The heatmap of log_10_ (relative abundance) of 48 HSV-1 L proteins in each subpopulation is shown. Data are the mean values from two biologically independent experiments. (B) Heatmap of the Pearson correlation coefficient between log_10_ (MFI of Venus) of each subpopulation and log_10_ (relative abundance) of each HSV-1 L protein. P-values for the correlation coefficient of all HSV-1 L proteins except Us8.5 shown in (B) were < 0.05.

### Specific Venus-Us11 protein expression levels are linked to progeny virus production

To analyze a relationship between expression levels of HSV-1 L proteins and progeny virus yields directly and quantitatively, HeLa cells were infected with rICP47/vUs11 at an MOI of 5, harvested at 4, 6, 8, 10, 12, and 24 h after infection, analyzed by flow cytometry and sorted into entire population (FSC singlet) or six subpopulations (f1 to f6) according to the levels of Venus-Us11 fluorescence intensity (S-Fig. 2). Virus titers in the entire population and each of the subpopulations were determined and virus titers per 10^4^ cells were estimated.

In agreement with previous reports ^4,6^, the kinetics of Venus-Us11 fluorescence intensity were similar to those of progeny virus titers (Fig. 3A) and Venus-Us11 fluorescence intensities had a high correlation with progeny virus titers at the entire population level (r=0.95) (Fig. 3B). We also calculated virtual titers per 10^4^ cells at each timepoint by summating virus titers in subpopulations, obtained by multiplying the estimated virus titer of each subpopulation of 10^4^ cells by the ratio of number of cells in the subpopulation to that in the entire population. A viral growth curve based on the virtual titers was almost identical to that based on actual titers in the entire population (Fig. 3C).

**Fig. 3.**
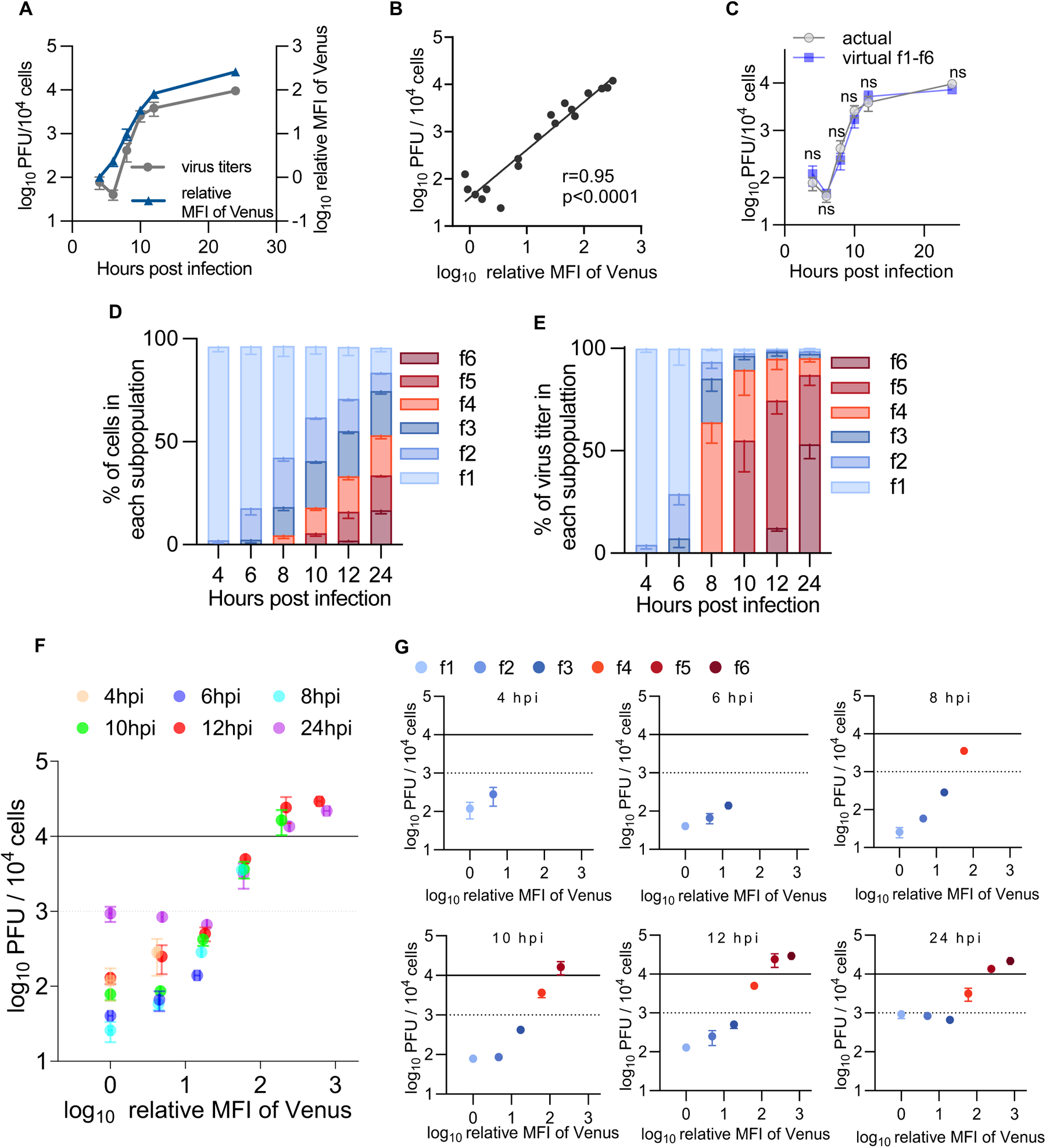
Quantitative analysis of a relationship between the expression levels of HSV-1 L proteins and progeny virus yields. (A to G) HeLa cells were infected with rICP47/vUs11 at an MOI of 5 and sorted as shown in S-Fig. 2. Sorted cells were sonicated and virus titers were determined by plaque assay using Vero cells. (A) Kinetics of mean fluorescent intensity (MFI) of Venus and progeny virus titers of the entire population. (B) Scatter plot of log_10_(relative MFI of Venus) vs log_10_(PFU/10^4^ cells) from panel A. (C) Kinetics of progeny virus titers of the entire population from panel A (actual) and sum of the virus titers produced by f1 to f6 subpopulation (virtual_f1-f6). The calculation method for virtual titers is described in the Materials and methods. (D) Proportion of cells in the indicated subpopulation account for the entire cell population at the indicated times after infection. (E) Proportion of virus titers produced by the f1 to f6 subpopulations account for virus titers produced by the entire population at the indicated times after infection. (F) Scatter plots of log_10_(relative MFI of Venus) vs log10(PFU/10^4^ cells). (G) Scatter plots of log_10_(relative MFI of Venus) vs log_10_(PFU/10^4^ cells) of the indicated subpopulations separated by time after infection from panel F. Each value is the mean ± standard error (A, C, D, E, F, and G) or individual value (B) of the results of three independent experiments. Statistical analysis was performed by unpaired Student’s *t*-test. ns, not significant (C). Solid and dashed lines indicate 1 PFU/cell and 0.1 PFU/cell, respectively (F and G).

At the entire population level, virus titers were decreased 6 h after infection due to an eclipse phase of infection and viral replication entered the productive phase between 6 and 8 h post-infection (Fig. 3A and C). These results indicated that infectious progeny virus production was detectable 8 h after infection in these experiments. The f4 subpopulation emerged at this timepoint (Fig. 3D, F and G, and S-Table 2). Although 4.1% of total cells were in the f4 subpopulation 8 h after infection (Fig. 3D, and S-Table 2), 55% of progeny infectious virus yields were produced by this subpopulation (Fig. 3E, and S-Table 2). The f5 and f6 subpopulations emerged 10 and 12 h after infection, respectively, when the growth rate had slowed (Fig. 3D, F and G, and S-Table 2). At 10 h after infection, 18% of total cells were in the f4 and f5 subpopulations (Fig. 3D, and S-Table 2) and these subpopulations produced 87% of progeny infectious virus yields (Fig. 3E, and S-Table 2). At 12 and 24 h post-infection, 33% and 53% of total cells were in the f4 to f6 subpopulations, respectively (Fig. 3D, and S-Table 2), and these subpopulations produced 95% of progeny infectious virus yields (Fig. 3E, and S-Table 2). Similarly, most progeny infectious virus yields were produced by the f4 to f6 subpopulations in HeLa cells using MOIs of 1 and 2.5 at 24 h post-infection and in other cells (Vero, U2Os and HaCaT cells, and human fetal foreskin fibroblasts (HFFF-2)) at MOIs 1 or 2.5 at 12 h post-infection, although the proportion of each subpopulation varied by different MOI and cell type (S-Figs. 3A to D and 4A to D, F to I, K to N and P to S). Notably, the viral growth curve from 8 h post-infection based on the virtual titers using only the data of the f4 to f6 subpopulations was almost identical to that based on the virtual titers using the data of the f1 to f6 populations or actual titers of the entire population (Fig. 4A and B). Similarly, virtual titers using only the data of the f4 to f6 subpopulations in HeLa cells at MOIs of 1 and 2.5 and in Vero, U2OS, and HaCaT cells, and HFFF-2 were also similar to virtual titers using the data of f1 to f6 populations or actual titers in the entire population (S-Figs. 3E to G, 4E, 4J, 4O, and 4T). These results indicated that subpopulations f4 to f6 have a predominant role in yielding progeny infectious viruses, whereas subpopulations f1 to f3 barely produced infectious virions. To obtain evidence to further support this conclusion, we analyzed virion morphogenesis by quantitating the number of virus particles at different morphogenetic stages in HSV-1-infected HeLa cells at an MOI of 5 for 8 (S-Fig. 5) or 24 h (Fig. 5) in each subpopulation f2 to f4 or f5, respectively, by electron microscopy. All virion types were barely detectable in infected cells from subpopulation f2 (Fig. 5 and S-Fig. 5). In subpopulation f3, although nuclear virions were obvious, cytoplasmic virions, and especially enveloped virions in the cytoplasm that were considered infectious, were barely detectable (Fig. 5 and S-Fig. 5). In contrast, enveloped virions were detected in the cytoplasm of 100% of infected cells in the f4 subpopulation at 8 h post-infection (S-Fig. 5) and these virions were detected in the cytoplasm of 70% and 100% of infected cells in the f4 and f5 subpopulations, respectively, at 24 h post-infection (Fig. 5). The proliferative profiles of enveloped virions in the cytoplasm (Fig. 5 and S-Fig. 5) were similar to those of infectious virus titers (S-Fig. 6A).

**Fig. 4.**
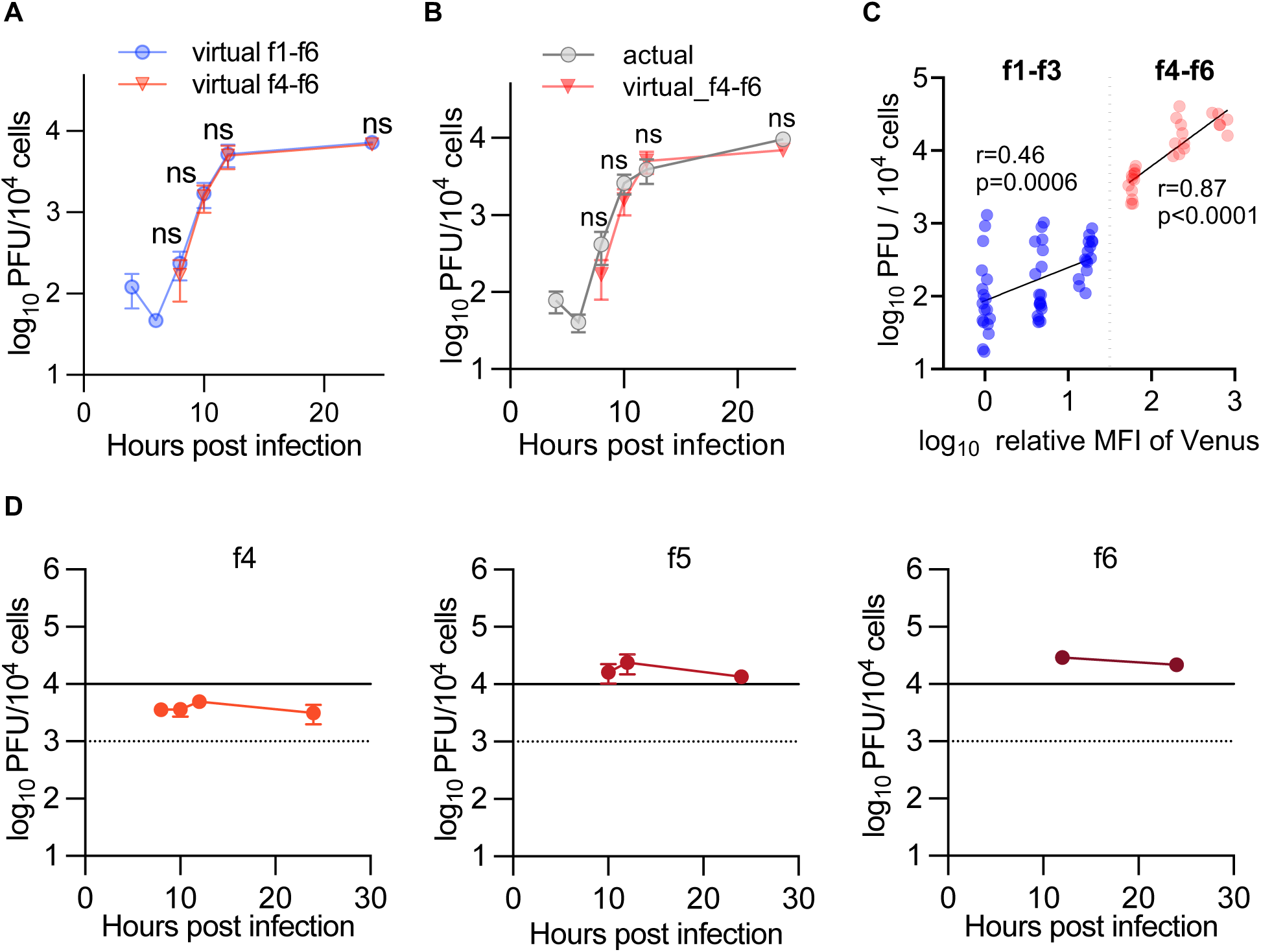
Cells expressing higher than specific levels of HSV-1 L proteins have a predominant role in yielding progeny infectious viruses. (A) Kinetics of the sum of the virus titers produced by f1 to f6 (virtual_f1-f6) or f4 to f6 (virtual_f4-f6) subpopulations. (B) Kinetics of progeny virus titers of the entire population (actual) and sum of the virus titers produced by f4 to f6 subpopulations (virtual_f4-f6). (C) Scatter plot of log_10_(relative MFI of Venus) vs log_10_(PFU/10^4^ cells) of subpopulations. (D) Progeny virus titers of the f4 to f6 subpopulations at the indicated times after infection. Each value is the mean ± standard error (A, B and D) or individual value (C) of the results of three independent experiments. Statistical analysis was performed by unpaired Student’s *t*-test. ns, not significant. All data are obtained from the experiment shown in Fig. 3.

**Fig. 5.**
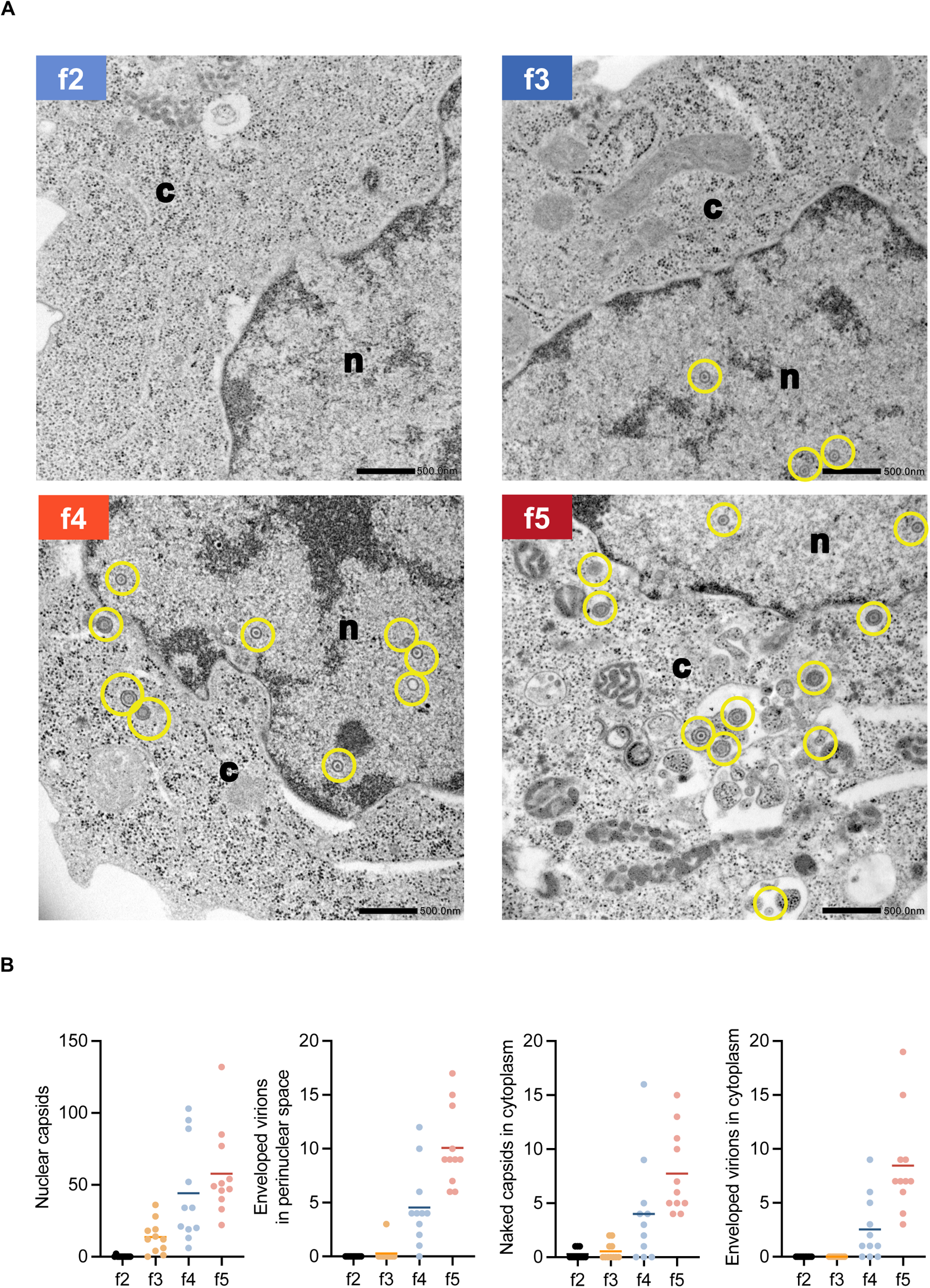
Electron microscopic analysis of cells in the f2 to f5 subpopulations. (A and B) HeLa cells were infected with rICP47/vUs11 at an MOI of 5 and sorted into four subpopulations (f2 to f5) by cell sorting 24 h after infection. Sorted cells were fixed, embedded, sectioned, stained, and examined by electron microscopy. (A) A transmission electron microscopy image of cells in the f2 to f5 subpopulations. n, nucleus; c, cytoplasm. Scale bars = 500 nm. (B) The numbers of nuclear virions, enveloped virions in the perinuclear space, naked capsids in the cytoplasm, and enveloped virions in the cytoplasm of 11 cells in the f2 to f5 subpopulations were quantitated. The horizontal bars indicate the means. Virions are marked in yellow.

**Fig. 6.**
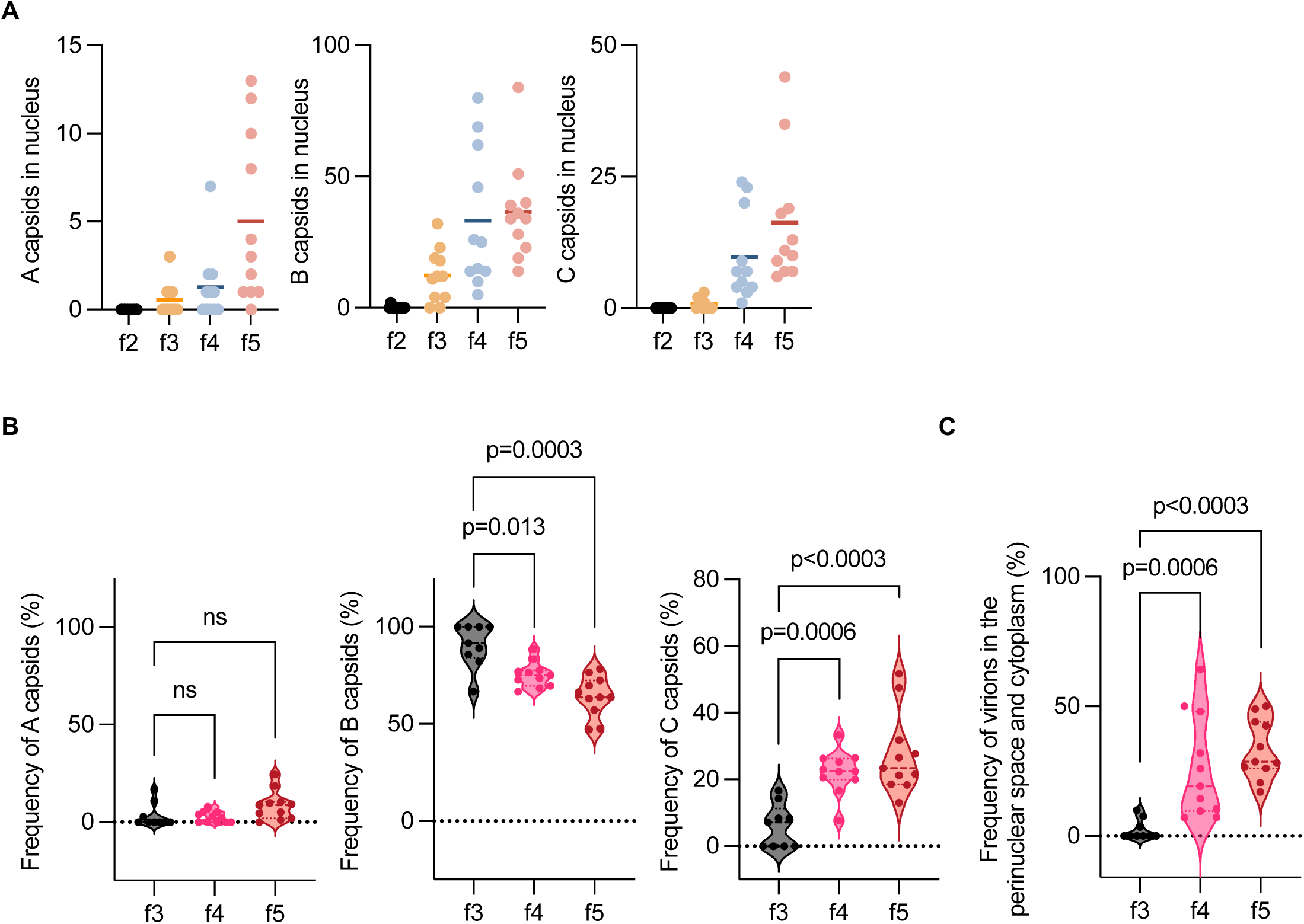
Frequencies of A, B, and C capsids in the nucleus of cells in the f2 to f5 subpopulations. (A) The numbers of A, B, and C capsids in the nucleus of 11 cells analyzed in Fig. 4 in the f2 to f5 subpopulations were quantitated. The horizontal bars indicate the means. Proportions of A, B, and C capsids to nuclear capsids (B), or virions in the perinuclear space and cytoplasm (C) of cells with more than two nuclear capsids (f3, n = 9; f4, n = 11; f5, n = 11). Data are presented as the median ± interquartile range (IQR). Statistical analysis was performed by the Mann-Whitney *U*-test, and P-values were adjusted by Bonferroni correction. ns, not significant.

We noted that, even at 4 and 6 h post-infection, the virus titers in f1 to f3 subpopulations were detectable at a maximum of 2.8 × 10^2^ PFU/10^4^ cells (Fig. 3G and S-Table 2). Taken together with the series of observations above (Figs. 3D and E, 4A and B, and 5, and S-Figs. 3, 4, 5, and 6A) indicating that infected cells in subpopulations f1 to f3 barely produced infectious virions, it was not likely that infectious virus titers in subpopulations f1 to f3 detected ranging from 2.6 × 10^1^ to 9.3 × 10^2^ PFU/10^4^ cells (Fig. 3G and S-Table 2) represented progeny virus yields of infected cells in f1 to f3 subpopulations. Cells in the f1 to f3 subpopulations were likely to be aborted infected cells and their frequencies 24 h after infection were similar to those reported previously in HeLa cells ^13,22^.

Collectively, these results together with the observation that Venus-Us11 fluorescence intensities reflected the expression levels of global HSV-1 L proteins indicated that certain levels of L protein expression were required for progeny virus production and a threshold of progeny virus production existed between Venus-Us11 protein expression levels in subpopulations f3 and f4. Notably, Venus-Us11 fluorescence intensities in subpopulations above the threshold (f4 to f6) correlated highly with progeny virus titers (r=0.87) (Fig. 4C). In contrast, a much lower correlation of coefficient (r=0.46) was observed in subpopulations below the threshold (Fig. 4C). Furthermore, virus titers in subpopulations f4 to f6 remained unchanged during HSV-1 infection (Fig. 4D and S-Fig. 6B) indicating L protein expression levels in subpopulations above the threshold were tightly correlated with virus titers independent of the timepoint after infection. Thus, infectious progeny virus production in the entire population over time depended on the proportion of these subpopulations in the entire population.

### Virion morphogenesis with a defect in the cleavage/packaging of HSV-1 DNA genome occurs in the subpopulation below the threshold for progeny virus production

The threshold for progeny virus production detected between Venus-Us11 protein expression levels in subpopulations f3 and f4 suggested a rate-limiting viral step for progeny virus production. Conceivably, this defect(s) might be observed in subpopulations below the threshold (f1 to f3) but not in subpopulations above the threshold (f4 to f6). To clarify this rate-limiting viral step, we compared levels of HSV-1 DNA genome replication and global viral mRNA expression in subpopulations f1 to f6 by quantitative PCR and RNA sequencing, respectively. Amounts of HSV-1 genome DNA increased in the f1 subpopulation and almost reached a plateau between subpopulations f2 and f3 (S-Fig. 7A). Similar proliferative profiles were observed for expressions of each of the 73 HSV-1 mRNAs tested (S-Fig. 7B). Thus, we did not detect any rate-limiting effects of HSV-1 genome replication or viral mRNA expression associated with the threshold for progeny virus production.

**Fig. 7.**
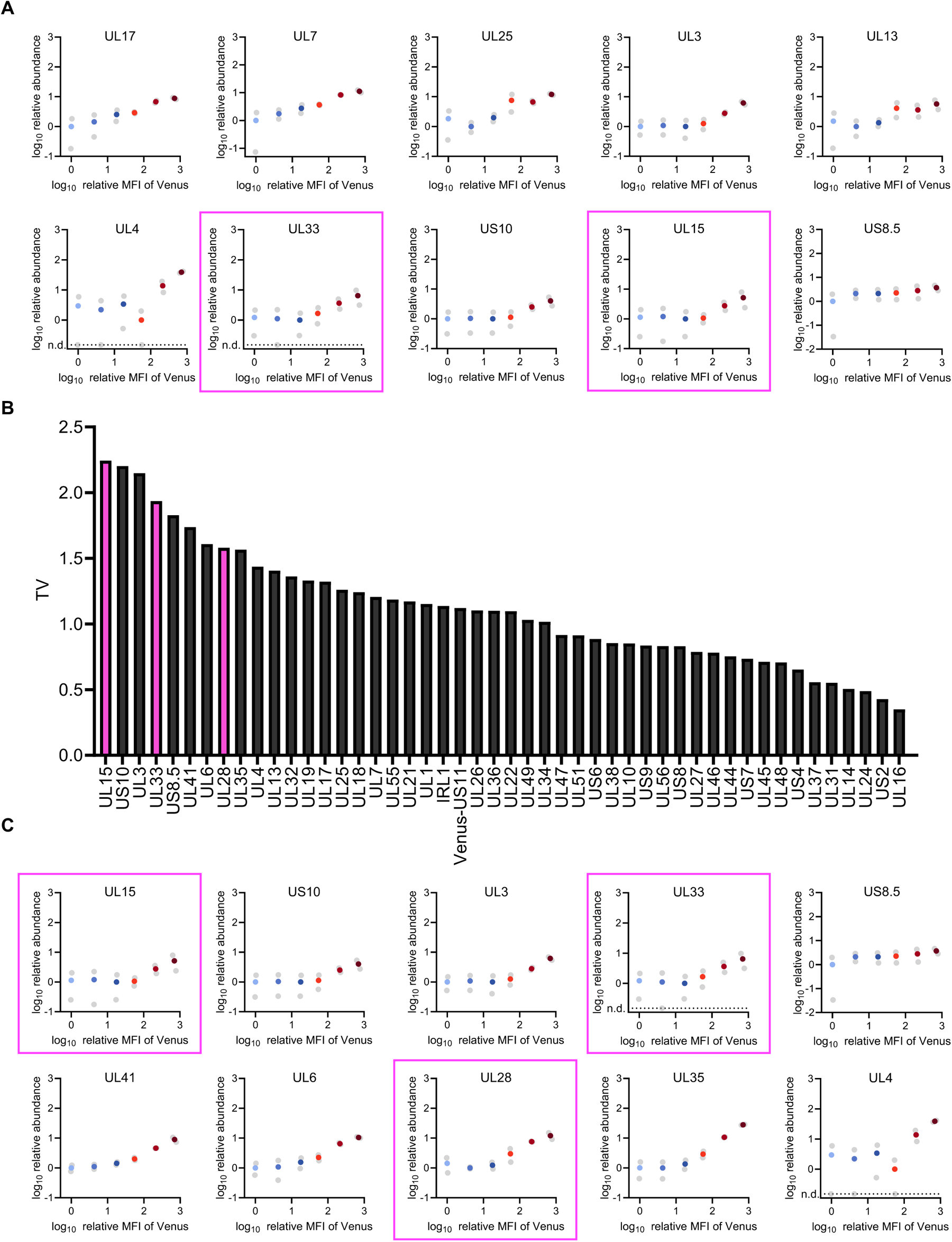
Unique protein expression profiles of a subset of HSV-1 L proteins. (A) Scatter plots of log_10_(relative MFI of Venus) vs log_10_(relative abundance) of L proteins (from S-Fig. 1B) of the bottom 10 HSV-1 L proteins with relatively low correlation coefficients between fluorescence intensity of Venus and their abundance (from Fig. 2B). Plots of terminase subunits are marked with magenta squares. (B) Threshold value (TV) of 48 L proteins (from S-Fig. 10). (C) Scatter plots of log_10_(relative MFI of Venus) vs log_10_(relative abundance) (from S-Fig. 1B) of the top 10 HSV-1 L proteins with highest TV (from S-Fig. 10). Data of terminase subunits are shown in magenta.

Next, we visualized virion morphogenesis by electron microscopy focusing on the A, B and C capsids in the nucleus (Fig. 6 and S-Fig. 8). The A and B capsids are incomplete structures resulting from problems in viral DNA genome retention in the capsids and packaging into the capsids, respectively ^23–25^. C capsids are mature capsids (nucleocapsids) containing viral DNA genomes reported to be selectively exported to the cytoplasm ^26–29^. B capsids accumulated aberrantly in subpopulation f3 and most (90.1%) nuclear capsids in this subpopulation were B capsids (Fig. 6 and S-Fig. 8). The frequency of B capsids in subpopulation f3 was significantly higher than in subpopulations f4 and f5 (Fig. 6B). In contrast, the frequency of C capsids in subpopulation f3 was significantly lower than in subpopulations f4 and f5 (Fig. 6B). In agreement with this and previous reports that C capsids are selectively exported to the cytoplasm ^26–29^, the frequency of virions in the perinuclear space and cytoplasm in subpopulation f3 was significantly lower than in subpopulations f4 and f5 (Fig. 6C). The frequency of A capsids in subpopulation f3 was comparable to those in subpopulations f4 and f5 (Fig. 6B). The frequencies of each type of nuclear capsid in subpopulations f4 and f5 were similar to those reported previously in wild-type HSV-1-infected HeLa cells ^30^. Furthermore, the aberrant accumulation of B capsids and lack of nuclear C capsids and virions in the perinuclear space and cytoplasm in subpopulation f3 were also observed at 8h post-infection (S-Figs. 5 and 9). These features of subpopulation f3 were similar to the phenotypes of HSV-mutants with defective viral DNA genome cleavage/packaging including those lacking the portal protein UL6, a terminase subunit (UL15, UL33 or UL28) or a minor capsid protein (UL17 or UL32) ^31–37^.

Collectively, these results suggested that the cleavage/packaging of HSV-1 DNA genome was specifically down-regulated in subpopulations below the threshold and that this viral step was a rate-limiting step for progeny virus production.

### A subset of HSV-1 L proteins including components of viral terminase have unique protein expression profiles

We compared global HSV-1 L protein abundance in each subpopulation by LC-MS/MS, and there was a strong correlation between the Venus-Us11 fluorescence intensities and relative abundance of most HSV-1 L proteins detected (Fig. 2B). Therefore, we focused on the abundance profiles of the bottom 10 HSV-1 L proteins with a relatively low correlation (Fig. 7A). The abundance profiles of several of these HSV-1 L proteins including UL25, UL3, UL4, UL33, Us10, and UL15 (Fig. 7A) differed from those of other HSV-1 L proteins whose abundance continuously increased in subpopulations f1 to f6 (S-Fig. 1B). The abundance of these L proteins in subpopulations f1 to f3 were very low and remained relatively constant, but were increased in subpopulations f4 to f6.

Next, to quantitatively evaluate the delayed increase in the abundance of L protein in the subpopulations, a four-parameter logistic regression was used for curve fitting of these profiles 24 h after infection (S-Fig. 10). The x-axis value at which the y-axis value increased by log_10_2 (2-fold increase in linear scale) from the starting point (x=0) for each curve was defined as the threshold value (TV) (S-Fig. 10 and Fig. 7B). Fig. 7C shows the top 10 proteins with the highest TV identified in this analysis. Of particular interest, these included all the subunits of HSV-1 terminase including UL15, UL33, UL28 and the HSV-1 portal protein UL6, all of which were reported to be required for viral DNA genome cleavage/packaging^31–34,36^. These results indicated that the expressions of a fraction of L proteins including all the subunits of HSV-1 terminase and the viral portal protein lacked in subpopulations f1 to f3, unlike that of other L proteins, suggesting the expressions of the L proteins are somehow downregulated in subpopulations below the threshold.

## DISCUSSION

Single-cell analyses have identified new subpopulations of virus-infected cells with similar viral gene expression profiles ^15–17^. However, the use of single-cell analyses to investigate both viral gene expression and progeny virus production is limited. Thus, single-cell analyses for both viral RNA synthesis and progeny virus production have been reported for some RNA viruses ^10,12,14^; a recent study reported that influenza virus transcription and progeny virus production were poorly correlated at the single cell level ^14^, and other studies did not focus on the relationship between viral transcription and virus progeny production at the single cell level ^10,12^. Furthermore, protein expression and virus progeny production at the single-cell or subpopulation level have not been investigated to date. This study established a system to fractionate HSV-1-infected cells based on global HSV-1 L protein expression levels and to titrate progeny virus yields in these fractions. This established system enabled us to clarify, for the first time, the direct and quantitative relationship between viral protein expression and progeny virus yields, and the clarified relationship indicated a threshold for HSV-1 L protein expression levels for progeny virus production. Such a threshold has not been reported by classical time course studies of most viruses at the entire population level by analyzing the relationship between viral gene expression and progeny virus yield. In contrast, once the levels of HSV-1 L protein expression exceeded the threshold, they were highly correlated with infectious progeny virus yields as reported by our (Fig. 3B) and other previous studies at the entire population level ^4,6^.

The threshold clarified in this study led us to identify the HSV-1 genome DNA cleavage/packaging step was a rate-limiting step for progeny virus production. Supporting this, MS analysis of the global expression of HSV-1 L proteins at the subpopulation level showed the expression of a subset of HSV-1 L proteins including all components of viral terminase and portal protein lacked in subpopulations below the threshold. Conceivably, an anti-viral strategy targeting this bottleneck step for HSV-1 production might block progeny virus production effectively. HSV-1 genome DNA cleavage/packaging, which is largely mediated by a specific viral enzyme HSV-1 terminase, is essential for viral replication ^31,33,34,36^, and most targets of antiviral drugs successfully developed to date are viral specific enzymes ^38^. Our study at the subpopulation level using this newly established system revealed an ideal target for the development of anti-viral drugs. Notably, the newly established system is simple and widely applicable for studies of the direct relationship between viral gene expression and progeny virus yield of many other DNA and RNA viruses, which might provide specific and conserved insights into mechanisms of progeny virus production in these viruses. It will be particularly interesting to determine whether the rate-limiting step identified in the progeny virus production of HSV-1 is conserved in adenoviruses, which seem to utilize similar DNA genome packaging systems ^39^.

It has long been recognized that much fewer virions are present in the cytoplasm of HSV-1-infected cells compared with the nucleus ^40–42^. Our conclusion that the HSV-1 genome DNA cleavage/packaging step is a rate-limiting step for progeny virus production is in agreement with these observations as well as earlier reports demonstrating C capsids are selectively exported to the cytoplasm ^26–29^. The ate-limiting step identified in this study also suggests a sophisticated strategy of efficient progeny virus production by HSV-1 without accumulating progeny nucleocapsids in the cytoplasm, which enables evasion from cytosolic sensors related to innate immune responses. In the cytoplasm, HSV-1 nucleocapsids are degraded by proteasomes ^43^, and viral DNAs are likely to be sensed by cytosolic sensors including cGAS, AIM2, DAI, and RNA polymerase III, which promote innate immune responses ^44–49^. Here we showed that, in subpopulation f3, which was likely slightly below the threshold, HSV-1 genome DNA replication had already reached a plateau and B capsid formation was easily observed although C capsid formation was barely detectable. In contrast, in subpopulation f4, which was likely slightly over the threshold, C capsids and all types of virions in the perinuclear space and the cytoplasm were evident. These observations suggested that downregulating the HSV-1 DNA cleavage/packaging might promote the accumulation of nucleocapsid components including capsids and viral genome DNAs in the nucleus as well as other viral proteins required for virion maturation at the NMs and in the cytoplasm at a level sufficient for efficient progeny virus production. Once the viral genome is packaged into a capsid, the following steps for virion maturation might immediately proceed without accumulating any excess nucleocapsids in the cytoplasm, decreasing the chance for HSV-1 to be sensed by innate immune responses in the cytoplasm. IFI16 and hnRNPA2B1 sense HSV-1 genome DNA and promote innate immune responses in the nucleus ^50–52^. However, replicated HSV-1 genome DNAs and nucleocapsids appear to accumulate in the nucleus of HSV-1-infected cells ^19,53,54^, suggesting this virus has evolved more effective evasion mechanisms against innate immune responses in the nucleus such that ICP0 counteracts IFI16 ^50^ and probably, against capsid degradation than those in the cytoplasm.

We noted that overall viral mRNA levels increased from the f1 subpopulation and almost reached a plateau between subpopulations f2 and f3 (S-Fig. 4B). In contrast, viral L protein levels continued to increase from the f1 to f6 subpopulations (S-Fig. 1B). Thus, there was a lack of correlation between viral L mRNA levels and viral L protein levels especially in fractions f4 to f6. In agreement with these observations, previous systematic studies quantifying mRNA and protein levels at the genomic scale reported the importance of multiple processes beyond the mRNA concentration that contributed to establishing the level of a protein ^55^. These processes include translation rates, translation rate modulation, modulation of a protein’s stability, protein synthesis delay, and protein transport ^55^. Interestingly, it was reported that HSV-1 VP22 promoted global viral protein expression ^56,57^, potentially by reducing the dependence of protein synthesis upon cellular ribosomal proteins to support translation during infection ^58^, and by promoting the translocation of viral L mRNAs to the cytoplasm ^59^. HSV-1 might evolve VP22 to promote global viral translation efficiency in specific circumstances such as in subfractions f4 to f6, in which amounts of viral mRNAs are saturated.

## MATERIALS AND METHODS

### Cells and Viruses

HeLa, U2OS, HFFF-2, HaCaT, and Vero cells were described previously ^20,60^. Wild-type HSV-1(F) was described previously ^61^.

### Generation of a recombinant virus

Recombinant virus YK410 (rICP47/vUs11) in which ICP47 and Us11 were tagged with TagRFP ^62^ and VenusA206K ^63^, respectively, were generated by two rounds of two-step Red-mediated mutagenesis using *Escherichia coli* GS1783 containing pYEbac102Cre ^61,64^, a full-length infectious HSV-1(F) clone, pBS-Venus-KanS ^20^, pFlag-TagRFP-KanS, and the primers listed in S-Table 1. The viruses used in this study were propagated and titrated in Vero cells.

### Plasmids

pFlag-TagRFP was constructed by amplifying the TagRFP open reading frame (ORF) to introduce a *Kpn*I site without changing the amino acid sequence by PCR from pTagRFP-N1 ^65^ and cloning it into pFlag-CMV2 (Sigma). pFlag-TagRFP-KanS, used in the two-step Red-mediated mutagenesis procedure, was constructed by amplifying the domain of pEP-KanS ^66^ carrying the I-*Sce*I site and the kanamycin resistance gene by PCR from pEP-KanS using the primers 5’-GCGGTACCGTGAACAACCACCACTTCAAAGGATGACGACGATAAGTAGGG-3’ and 5’-GCGGTACCCTCCATGTACAGCTTCATGTCAACCAATTAACCAATTCTGATTAG-3’ and cloning it into the *Kpn*I site of pFlag-TagRFP. pGEX-ICP47 was constructed by amplifying the ICP47 ORF by PCR from the HSV-1(F) genome and cloning it into pGEX-4T-1.

### Antibodies

Commercial antibodies used in this study included mouse monoclonal antibody to α-tubulin (DM1A; Sigma), and rabbit monoclonal antibodies to GFP (598; MBL) and TagRFP (AB233; Evrogen). Rabbit polyclonal antibodies to US11 were described previously ^41^. To generate a rabbit polyclonal antibody to ICP47, a rabbit was immunized, according to the standard protocol at MBL, with GST-ICP47 expressed in *E. coli* and purified as described previously ^20^. Serum from the immunized rabbit was used as the anti-ICP47 polyclonal antibody.

### Immunoblotting

Immunoblotting was performed as described previously ^67^.

### Flow cytometry

HeLa cells infected with HSV-1(F) or YK410 (rICP47/vUs11) were washed with phosphate buffered saline (PBS) and detached with 0.25% trypsin/EDTA solution (Wako). Then, the cells were suspended in PBS containing 2% FCS, filtered through a 35-μm pore-cell strainer (#352235, Corning) and subjected to FACS analysis with BD FACS Melody (Becton Dickinson). The data were analyzed with BD FACSChorus (Becton Dickinson) software or FlowJo 10.8.1 software (Becton Dickinson).

### Determination of viral titer in cells fractionated by cell sorting

HeLa cells were infected with YK410 (rICP47/vUs11) at an MOI of 5. At the indicated times after infection, cells were detached and suspended as described above. Then, 1.5 × 10^4^ to 3 × 10^4^ cells in the f1 to f6 subpopulations, and the entire cell population (FSC-singlet), were sorted into medium 199 containing 1% FCS by FACS Melody (Becton Dickinson). The sorted cells were freeze-thawed once, sonicated, and virus titers were determined by plaque assay using Vero cells. The average virus titer per single cell was obtained by dividing the obtained virus titer by the number of sorted cells. Data are presented as the PFU per 10^4^ cells.

### Calculation of the proportion of virus titers in each subpopulation and the virtual titer

PFU_fi_, which represents the viral titer of each subpopulation fi (i = 1 to 6), was obtained using the following equation:

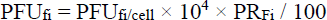

where PFU_fi/cell_ is the average viral titer per single cell belonging to the fi subpopulation (from Fig. 3F and G) and PR_fi_ is the proportion (%) of cells in the fi fraction in the entire population (from Fig. 3D). The virtual titer (VT), which represents the sum of PFU_fi_ was obtained as follows:

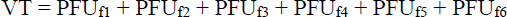

% of PFU_Pi_, which represents the proportion of viral titers in the fi subpopulation accounted for in the entire population (Fig. 3E) was obtained using the following equation:

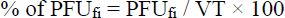

### LC-MS/MS analysis of cells fractionated by cell sorting

HeLa cells were infected with rICP47/vUs11 at an MOI of 5 for 24 h. Then, cells were detached and suspended as described above. Next, 10^5^ cells in the f1 to f6 subpopulation were sorted into PBS by BD FACS Melody (Becton Dickinson), pelleted by centrifugation, and lysed with lysis buffer (0.1 M of Tris-HCl (pH 8.0), 1% SDS). To remove SDS from samples, we used the methanol-chloroform protein precipitation method. Briefly, we added 4 volumes of methanol, 1 volume of chloroform, and 3 volumes of water to the eluted sample and mixed thoroughly. The samples were centrifuged at 15,000 rpm for 10 min, the water phase was carefully removed, and then 4 volumes of methanol were added to the samples. The samples were centrifuged at 15,000 rpm for 10 min, and the supernatant was removed. The pellet was washed once with 100% ice-cold acetone. The precipitated protein was re-dissolved in guanidine hydrochloride, reduced with TCEP, alkylated with iodoacetamide, and digested with lysyl endopeptidase and trypsin. The resulting digested peptides were analyzed using an Evosep One LC system (EVOSEP) connected to a Q-Exactive HF-X mass spectrometer (Thermo) with a Dream spray tip (AMR) and a 15 cm × 150 μm column packed with 1.9-μm C18-beads (Evosep). The mobile phases were comprised of 0.1% FA as solution A and 0.1% FA/99.9% ACN as solution B. The analysis was performed in the data-dependent acquisition mode, where the top 25 recorded mass spectrometry spectra between 380 and 1500 m/z were selected. Survey scans were acquired at a resolution of 60,000 at m/z 200, and the tandem mass spectrometry (MS/MS) resolution was set to 15,000 at m/z 200. All MS/MS spectra were searched against the protein sequences of the HSV protein database and human Swiss-Prot database using Proteome Discoverer 2.2 (Thermo) with the SEQUEST search engine, and the result of HSV proteins was extracted. The false discovery rate (FDR) was set to 1% on peptide spectrum match (PSM).

### Data processing of LC-MS/MS data

Normalization was performed on quantitative values using the method built into the Discoverer 2.2 software. Briefly, the Total Peptide Amount mode was used to normalize between samples, and the On All Average mode was used to normalize between identified proteins. The subsequent analysis was performed using only the extracted data for HSV proteins. The normalization method in each mode was as follows. The Total Peptide Amount mode sums the peptide group abundances for each sample and determines the maximum sum for all files. The normalization factor is the factor of the sum of the samples and the maximum sum in all files. The On All Average mode aggregates all the abundance or normalized abundance values per sample and scales the abundance values of each sample so that the average of all samples is 100. When a peptide was not detected in four technical replicates from one sample of each subpopulation, the protein was considered undetectable in that subpopulation. When a peptide was not detected in a replicate of each subpopulation, the missing value was complemented with the mean value of abundances in technical replicates where the peptide was detected. The mean value of abundances from four technical replicates were used as the value of a single experiment. The relative abundance of each HSV-1 L protein was calculated relative to the subpopulation with the lowest mean abundance for two biologically independent experiments as 1.

### RNA-sequencing in cells fractionated by cell sorting

HeLa cells were infected with YK410 (rICP47/vUs11) at an MOI of 5 for 24 h. Then, cells were detached and suspended as described above. Next, 5 × 10^5^ cells in the f1 to f6 subpopulations were sorted into PBS by BD FACS Melody (Becton Dickinson). After centrifugation, total RNA from fractionated cells was isolated with a High Pure RNA Isolation kit (Roche). (i) Each cDNA was generated using a Clontech SMART-Seq HT Kit (Takara Clontech, Mountain View, CA, USA), and each library was prepared using a Nextera XT DNA Library Prep Kit (Illumina, San Diego, USA). Sequencing was performed on the DNBSEQ-G400 platform in the 100+100-base paired-end mode. (ii) Generated reads were a mixture of human reads and HSV-1 reads. For human data, generated reads were mapped to the human (hg19) reference genome using TopHat v2.1.1 combined with Bowtie2 ver. 2.2.8 and SAMtools ver. 0.1.18. For HSV-1, generated reads were mapped to the HSV-1 (GenBank: GU734771.1) reference genome using HISAT2 v2.1.0. (iii) The quantification to obtain the read counts of each gene from HSV-1 was performed using featureCounts of subread-2.0.0 with option -M --fraction. With the length data of each gene (another output of the featureCounts), fragments per kilobase of exon per million mapped fragments (FPKMs) were calculated according to the definition. The FPKM of HSV-1 genes was multiplied by (total number of mapped reads of the HSV-1 genome)/{(total number of mapped reads of the human genome) + (total number of mapped reads of the HSV-1 genome)}, normalized by the amount relative to the value of the f1 fraction.

### Quantification of HSV-1 DNA in cells fractionated by cell sorting

HeLa cells were infected with YK410 (rICP47/vUs11) at an MOI of 5 for 24 h. Then, cells were detached and suspended as described above. Next, 5 × 10^4^ cells in the f1 to f6 subpopulations were sorted into PBS by BD FACS Melody (Becton Dickinson). After centrifugation, total DNA from fractionated cells was isolated with NucleoSpin Tissue XS (TAKARA) according to the manufacturer’s instructions. The amount of HSV-1 DNA was quantified using the Universal Probe Library (Roche) with TaqMan Master (Roche) and the LightCycler 96 System (Roche) according to the manufacturer’s instructions. Primers and probes were designed using probe finder software within the region encoding ICP27 in the HSV-1 genome, and within the region encoding 18S rRNA in the human genome. The primer and probe sequences for ICP27 (for HSV-1 DNA) were 5′-TCCGACAGCGATCTGGAC-3′, 5′-TCCGACGAGGAACACTCC-3′, and Universal ProbeLibrary probe 56, and for 18S rRNA (for human DNA) the sequences were 5′-GCAATTATTCCCCATGAACG-3′, 5′-GGGACTTAATCAACGCAAGC-3′, and Universal ProbeLibrary probe 48. The ΔCt was calculated by subtracting the Ct value of HSV-1 DNA by the Ct value of human DNA. The relative amount of HSV-1 DNA to human DNA (2^-ΔCt^) was normalized relative to the value of the f1 subpopulation.

### Electron microscopic analysis of cells fractionated by cell sorting

HeLa cells were infected with YK410 (rICP47/vUs11) at an MOI of 5 for 8 h or 24 h. Then, cells were detached and suspended as described above. Next, 5 × 10^5^ to 1 × 10^6^ cells in the f2 to f4 (8 h) or f2 to f5 (24 h) subpopulations were sorted by BD FACS Melody (Becton Dickinson). After centrifugation, the pelleted cells were examined by ultrathin-section electron microscopy as described previously ^68^ except using JEM-1400 Flash microscope (JEOL).

### Statistical analysis

The unpaired Student’s *t*-test was used to compare two groups. One-way ANOVA followed by the Tukey multiple comparisons test, or Mann-Whitney *U-*test followed by Bonferroni correction were used for multiple comparisons. A P-value < 0.05 was considered statistically significant. For the statistical comparison of viral titers, data were converted to common logarithms (log_10_). Pearson correlation coefficients (r) and p-values were calculated on log10 transformed data. GraphPad Prism 10 (GraphPad Software) was used to perform statistical analyses.

### Classification of HSV-1 genes

HSV-1 genes were classified into IE, E, and L genes as described previously ^19^.

### Curve fitting

To describe the abundance profiles of HSV-1 L proteins, we mainly used the following mathematical model developed by dose response modeling ^69^.

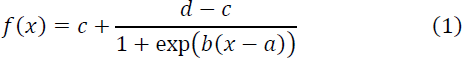

The variable *f*(*x*) is the abundance profiles of HSV-1 L proteins. The parameter *a* is the inflection point of the curve for this model, that is, the point where a change in acceleration in the curve occurs. The parameter *b* is a slope parameter and the parameters *c* and *d* are the lower and upper horizontal asymptotes or limits, respectively.

Fitting was performed by drc package in R, which estimates the parameters with nonlinear least squares under the assumption of normally distributed response values ^70^.

## ACKNOWLEDGEMENTS

We thank Risa Abe, Tohru Ikegami and Yui Muto for their excellent technical assistance. We acknowledge the NGS core facility of the Genome Information Research Center at the Research Institute for Microbial Diseases of Osaka University for the support in RNA sequencing and data analysis.

## Funding

This study was supported by Grants for Scientific Research and Grant-in-Aid for Scientific Research (S) (20H05692) from the Japan Society for the Promotion of Science (JSPS), grants for Scientific Research on Innovative Areas (21H00338, 21H00417, 22H04803) and a grant for Transformative Research Areas (22H05584) from the Ministry of Education, Culture, Science, Sports and Technology of Japan, PRESTO (JPMJPR22R5), SPRING (JPMJSP2018), MIRAI (JPMJMI22G1) and Moonshot R&D (JPMJM2021, JPMJMS2025) from Japan Science and Technology Agency (JST), grants (JP20wm0125002, JP22fk0108640, JP22gm1610008, JP223fa627001, JP23wm0225031, JP23wm0225035) from the Japan Agency for Medical Research and Development (AMED), grants from the International Joint Research Project of the Institute of Medical Science, the University of Tokyo, and grants from the Takeda Science Foundation, the Uehara Memorial Foundation and the Mitsubishi Foundation.

## Author contributions

Conceptualization: MN, YM, YK; Methodology: MN, YM, YK; Investigation: MN, YM, KT, FM, HK, TN, SA; Resources: MN, YM, KT, NK, AK; Formal analysis: MN, YM, RY, TN, HP, YK, SI; Visualization: MN, YM, RY, TN, HP, YK, SI; Funding acquisition: YM, SI, AK, YK; Writing – original draft: MN, YM, YK; Supervision: YK

## Competing interests

The authors declare that they have no competing interests.

## Data and materials availability

All data needed to evaluate the conclusions in the paper are present in the paper and/or the Supplementary Materials. The mass spectrometry data were deposited in the Japan ProteOme STandard Repository (jPOST) under the ID JPST002328. RNA-seq data have been deposited to Gene Expression Omnibus (GEO) with accession number GSE240449.

## Notes

### Competing Interest Statement

The authors have declared no competing interest.

### Summary of Updates

Figures, Supplementary Figures, and the manuscript are revised.

