## Supplemental Figures and Tables for "Direct Relationship between Protein Expression and Progeny Yield of Herpes Simplex Virus 1 Unveils a Rate-limiting Step for Virus Production"

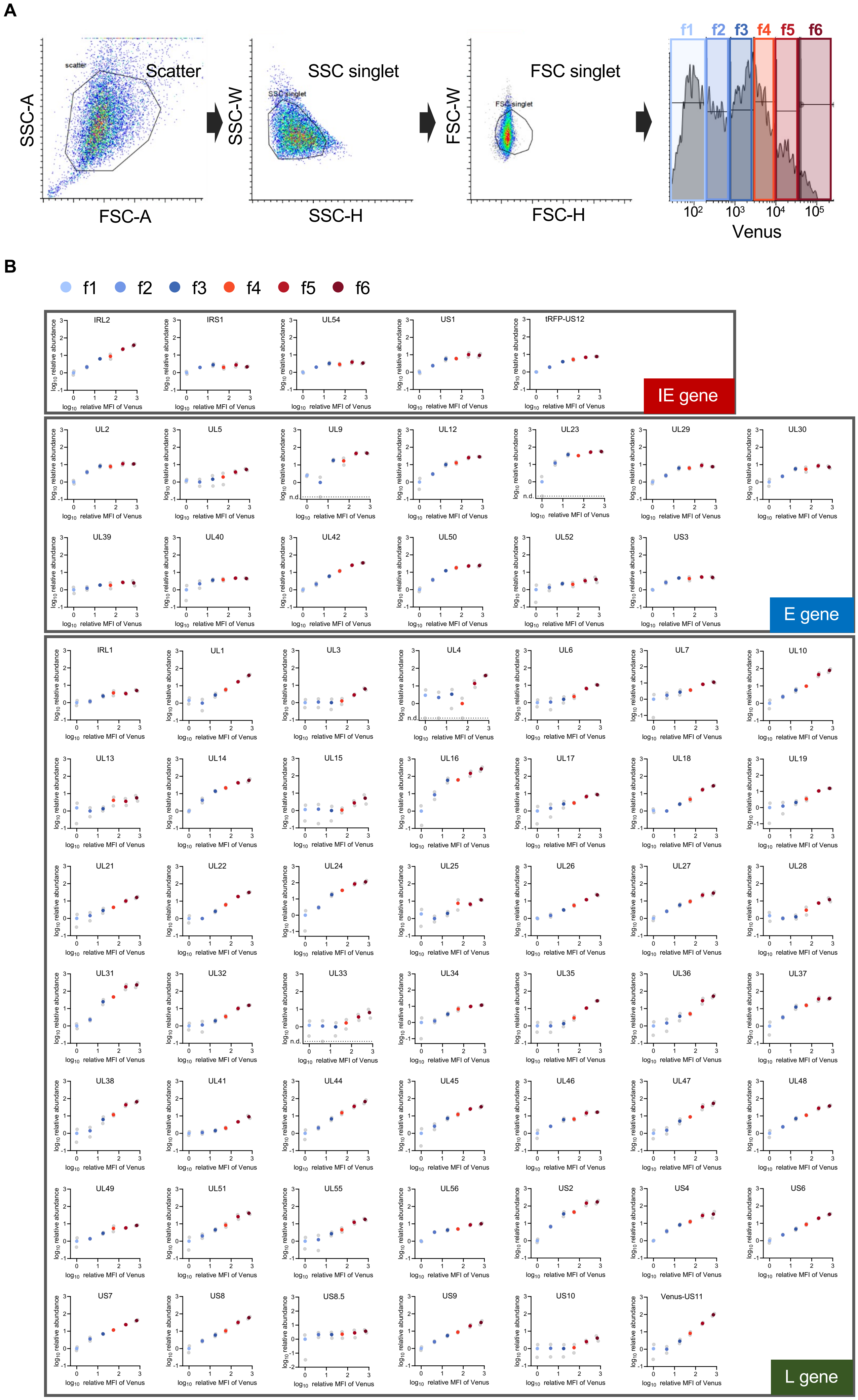


*(Legend on next page)*

**Supplementary Fig. 1 M. Nobe and Y. Maruzuru et al.**

**Supplementary Fig. 1. Abundance of HSV-1 L proteins in each subpopulation of rICP47/vUs11 infected cells.** (A) Gating strategy for the LC-MS/MS analysis of HeLa cells infected with rICP47/vUs11 at an MOI of 5 for 24 h. (B) Scatter plots of $log_{10}$(relative MFI of Venus) vs $log_{10}$(relative abundances) of the 5 indicated HSV-1 IE proteins, 13 indicated E proteins, and 48 indicated L proteins. Each value is the mean of two biologically independent experiments. Individual value from two independent experiments is plotted as a gray circle.

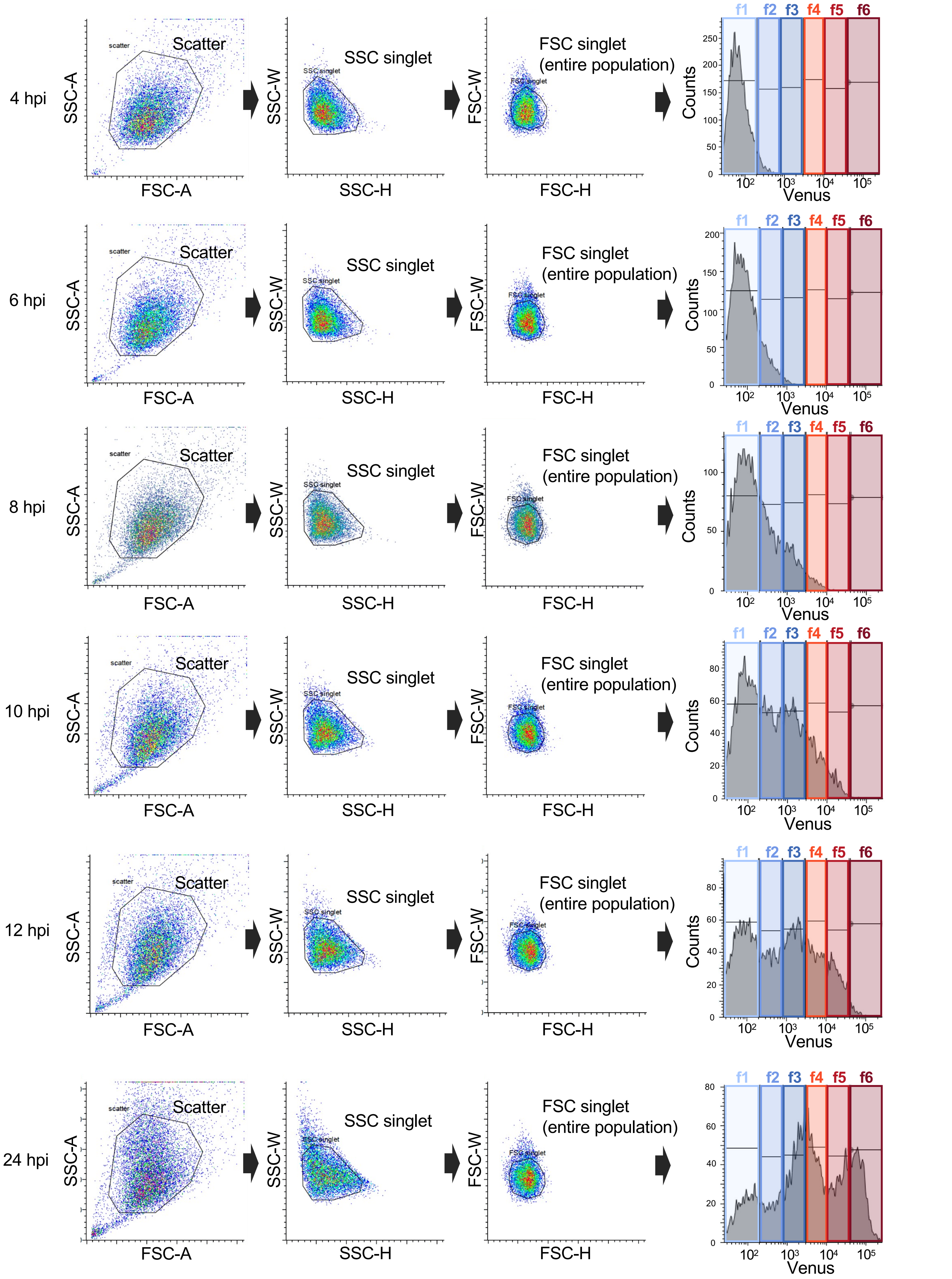


**Supplementary Fig. 2. Gating strategy for the experiments shown in Fig. 3.** HeLa cells were infected with rICP47/vUs11 at an MOI of 5, analyzed by flow cytometry, and sorted into an entire cell population (FSC singlet) or f1 to f6 subpopulations by cell sorting at the indicated times after infection. Sorted cells were analyzed as shown in Fig 3.

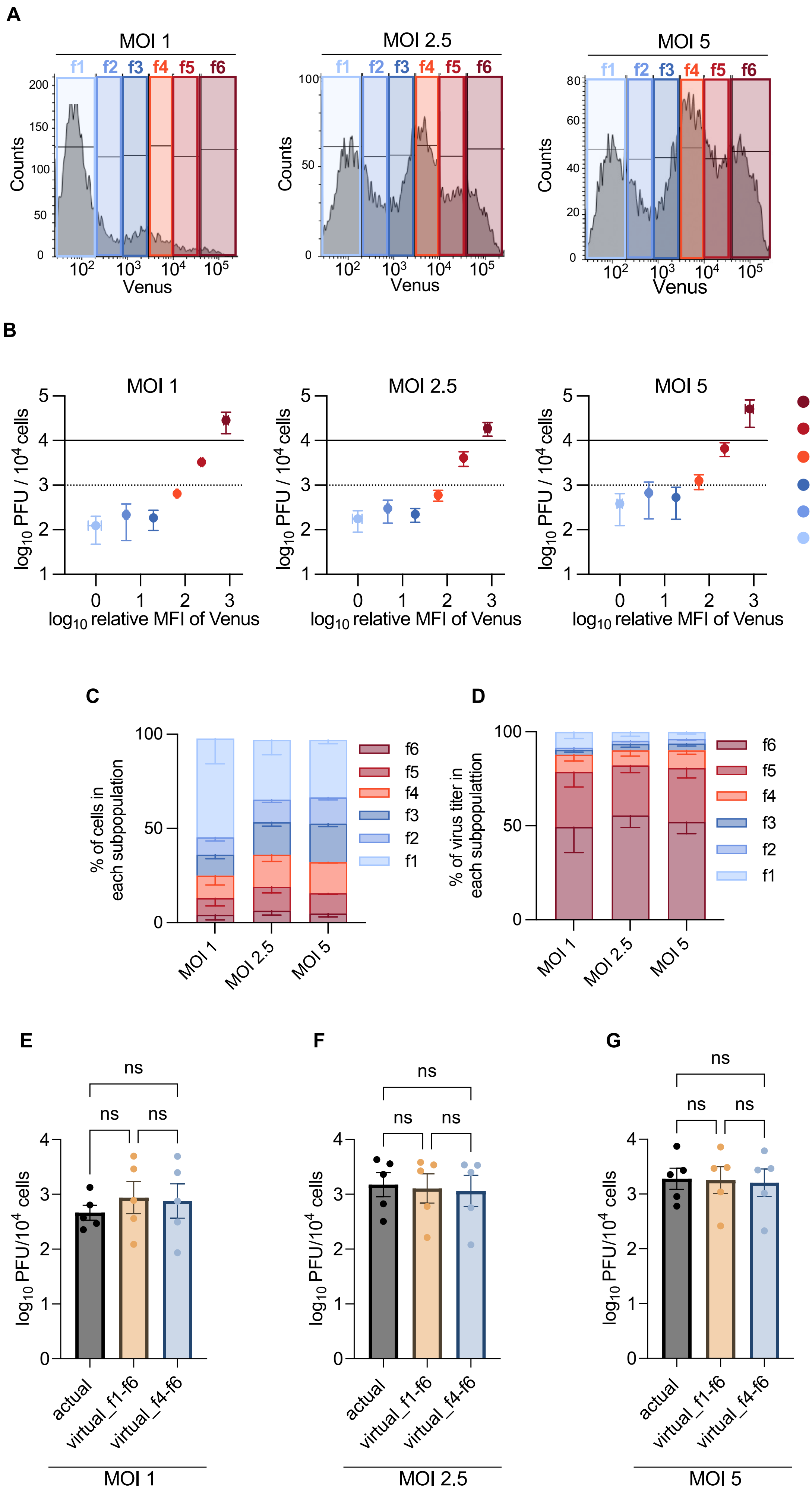


*(Legend on next page)*

**Supplementary Fig. 3 M. Nobe and Y. Maruzuru et al.**

**Supplementary Fig. 3. Quantitative analysis of the relationship between the expression levels of HSV-1 L proteins and progeny virus yields at different MOIs.** (A to G) HeLa cells infected with rICP47/vUs11 at the indicated MOI for 24 h, were sorted into f1 to f6 subpopulations (A), sonicated and virus titers were determined by plaque assay using Vero cells. (B) Scatter plot of $\log_{10}$(relative MFI of Venus) vs $\log_{10}$(PFU/$10^4$ cells). (C) Proportion of cells in the indicated subpopulations account for the entire cell population at the indicated MOIs. (D) Proportion of virus titers produced by the f1 to f6 subpopulations account for virus titers produced by the entire population at the indicated times after infection. (E to G) Progeny virus titers of the entire population (actual) and sum of the virus titers produced by f1 to f6 (virtual_f1-f6) or f4 to f6 (virtual_f4-f6) subpopulations at the indicated MOIs. The data are representative of five independent experiments (A). Each value is the mean ± standard error of the results of five independent experiments (B to G). Statistical analysis was performed by one-way ANOVA followed by the Tukey test (E to G). n.s., not significant

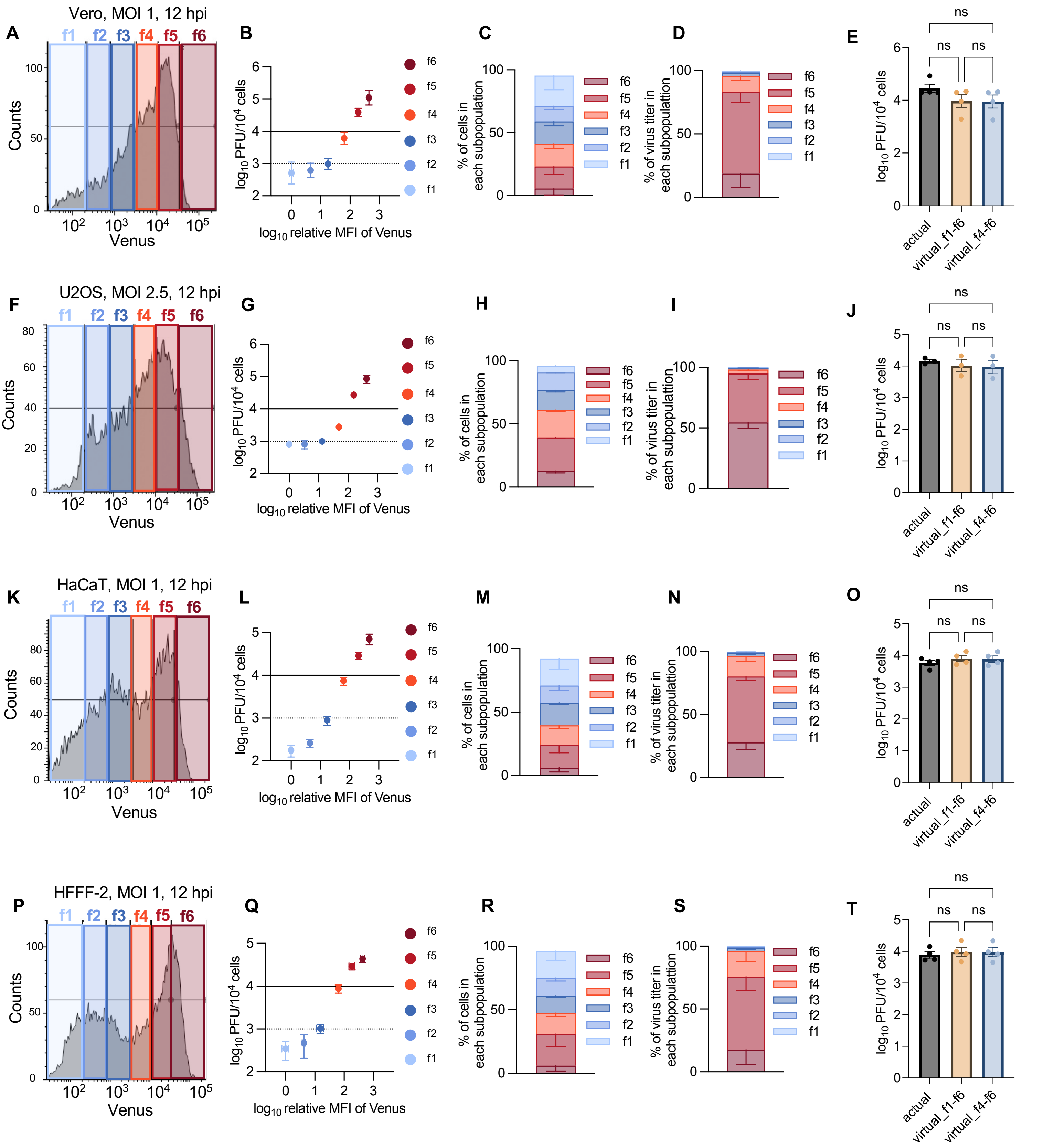


**Supplementary Fig. 4. Quantitative analysis of the relationship between the expression levels of HSV-1 L proteins and progeny virus yields using different cell lines.** Vero (A to E), U2OS (F to J), HaCaT (K to O), or HFFF-2 (P to T) cells infected with rICP47/vUs11 at an MOI of 1 (A to E, and K to T) or 2.5 (F to J) for 12 h, were sorted into f1 to f6 subpopulations (A, F, K, and P), sonicated, and virus titers were determined by plaque assay using Vero cells. (B, G, L, and Q) Scatter plots of $\log_{10}$(relative MFI of Venus) vs $\log_{10}$(PFU/$10^4$ cells) of each subpopulation. (C, H, M, and R) Proportion of cells in the f1 to f6 subpopulations account for the entire cell population. (D, I, N, and S) Proportion of virus titers produced by each subpopulation account for the virus titers produced by the entire cell population. (E, J, O, and T) Progeny virus titers of the entire population (actual) and sum of the virus titers produced by f1 to f6 or f4 to f6 subpopulations. The data are representative of three (F) or four (A, K, and P) independent experiments. Each value is the mean ± standard error of the results of three (G to J) or four (B to E, L to O, and Q to T) independent experiments. Solid and dashed lines indicate 1 PFU/cell and 0.1 PFU/cell, respectively (B, G, L and Q). Statistical analysis was performed by one-way ANOVA followed by the Tukey test. n.s., not significant (E, J, O, and T).

**Supplementary Fig. 4 M. Nobe and Y. Maruzuru et al.**

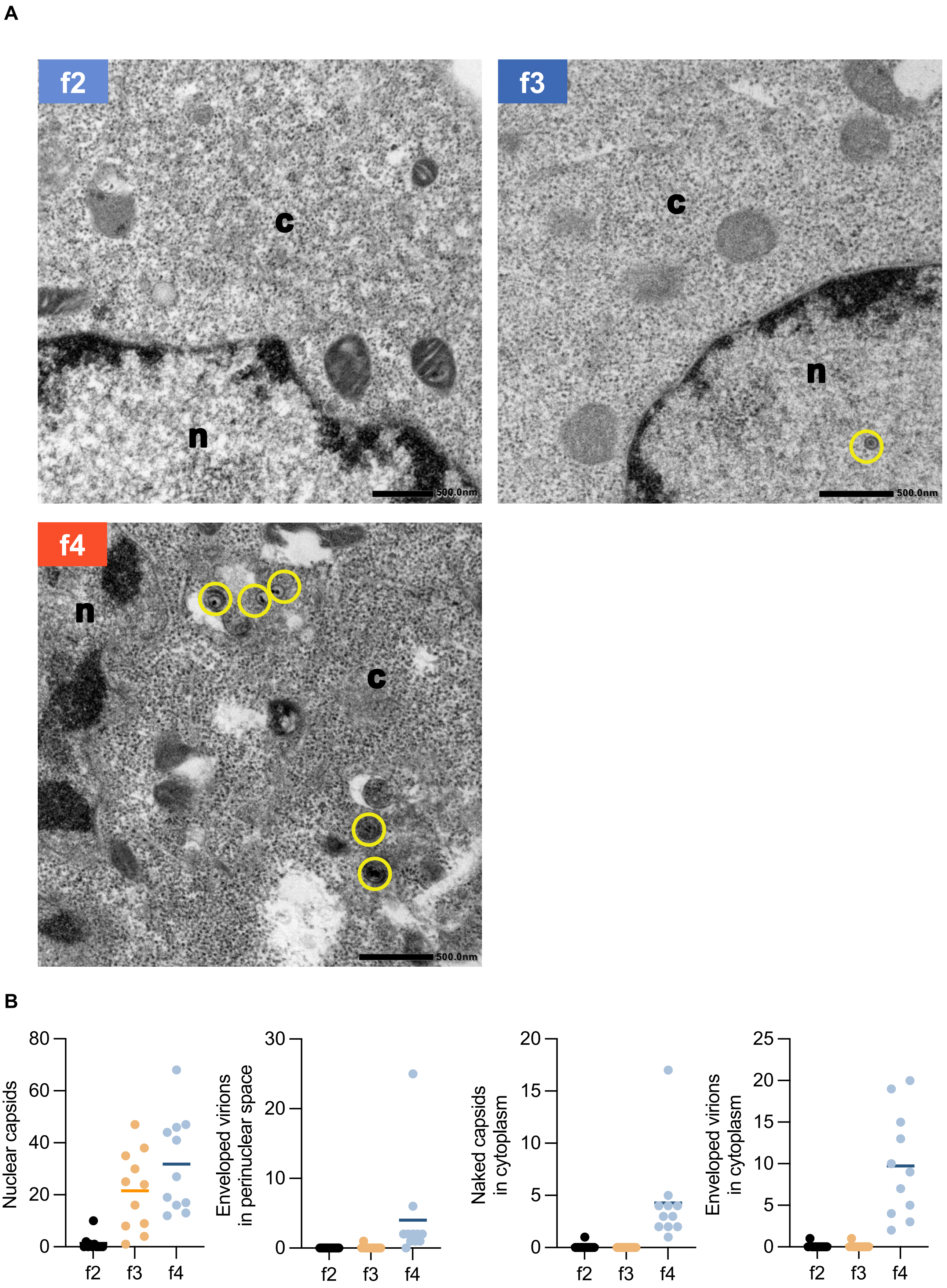


**Supplementary Fig. 5. Electron microscopic analysis of cells in the f2 to f4 subpopulations at 8 h after infection.** (A and B) HeLa cells were infected with rICP47/vUs11 at an MOI of 5 and sorted into three subpopulations (f2 to f4) by cell sorting 8 h after infection. Sorted cells were fixed, embedded, sectioned, stained, and examined by electron microscopy. (A) A transmission electron microscopy image of cells in the f2 to f4 subpopulations. n, nucleus; c, cytoplasm. Scale bar = 500 nm. (B) The numbers of nuclear virions, enveloped virions in the perinuclear space, naked capsids in the cytoplasm, and enveloped virions in the cytoplasm of 11 cells in the f2 to f4 subpopulations were quantitated. The horizontal bars indicate the means. Enveloped virions and naked capsids are marked in yellow.

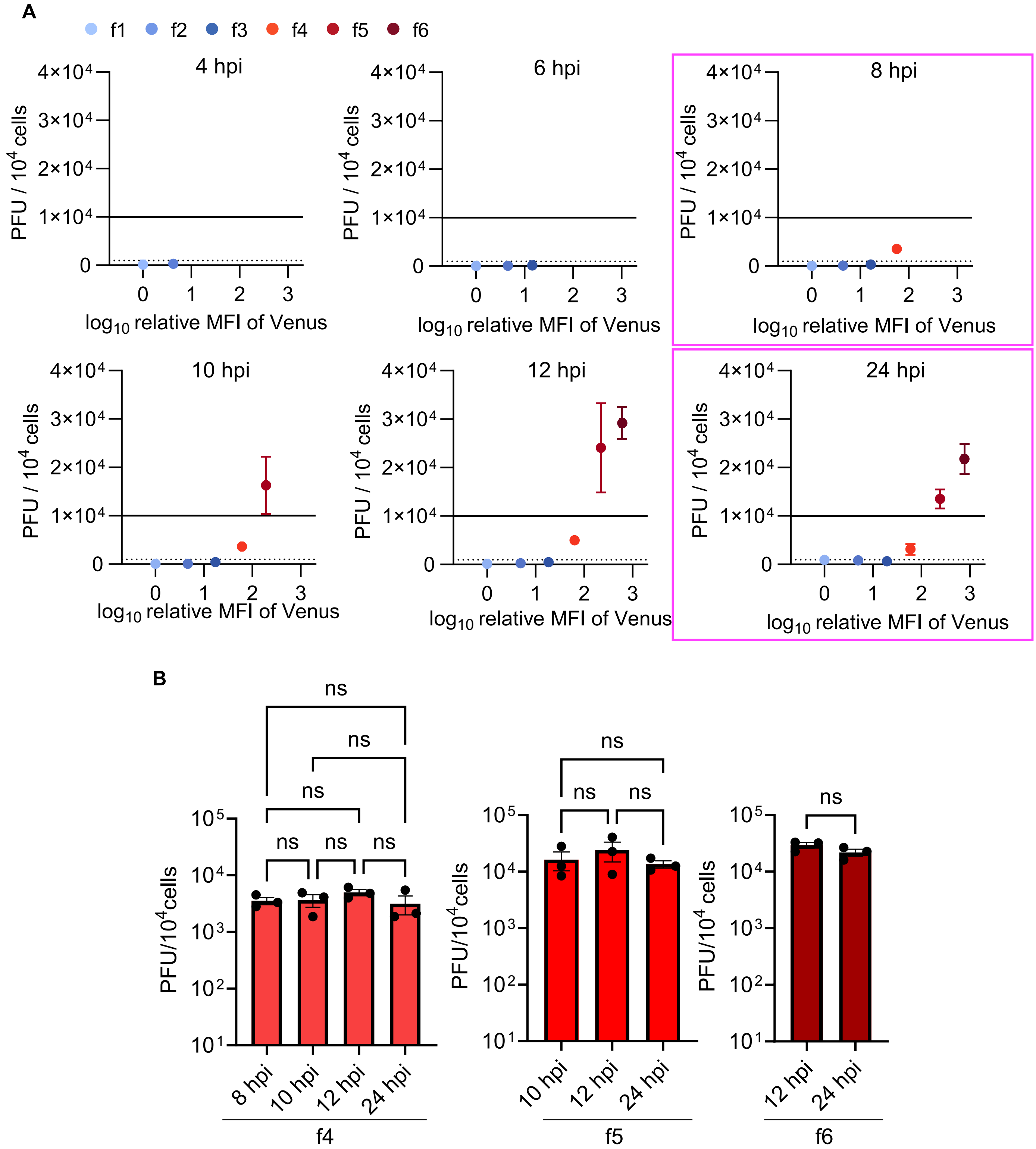


**Supplementary Fig. 6. F4 to f6 subpopulations have a predominant role in yielding progeny infectious viruses.** (A) Scatter plots of $log_{10}$(relative MFI of Venus) vs PFU/$10^4$ cells of the indicated subpopulation separated by time after infection from Fig. 3 panel G. Scatter plots under the same conditions of electron microscopic analysis performed in Fig. 5 and S-Fig. 5 are shown in magenta. (B) Progeny virus titers of the f4 to f6 subpopulations at the indicated times after infection from Fig. 4 panel B. Statistical analysis was performed by one-way ANOVA followed by the Tukey test. n.s., not significant.

A

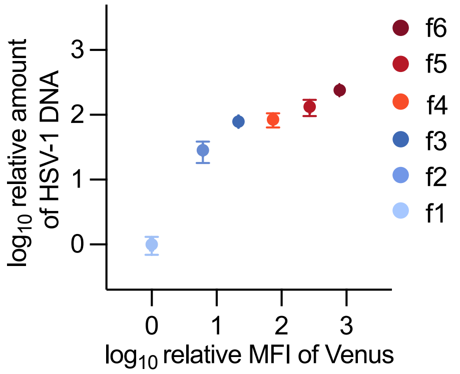


B

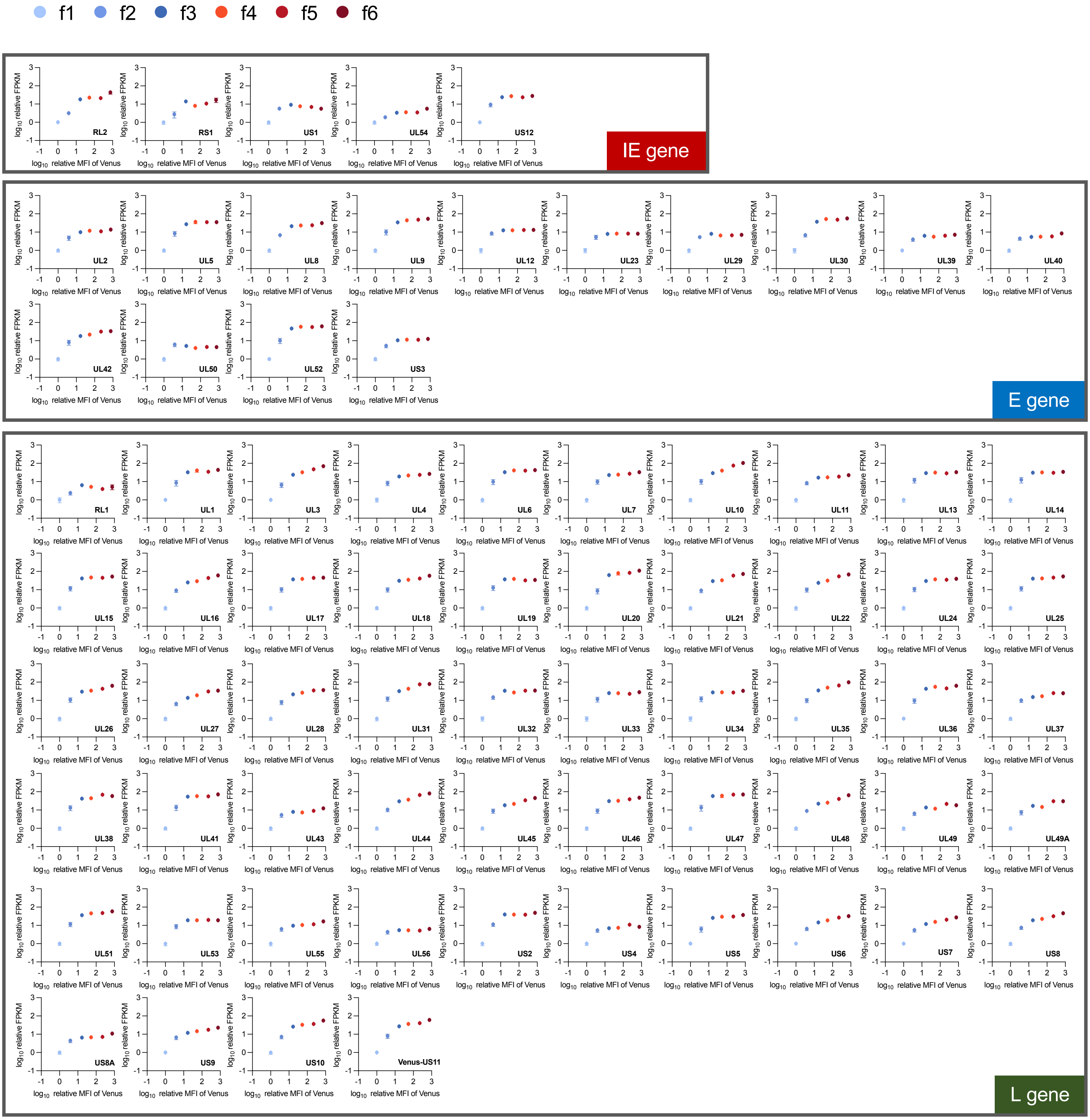


*(Legend on next page)*

**Supplementary Fig. 7 M. Nobe and Y. Maruzuru et al.**

**Supplementary Fig. 7. Amount of HSV-1 DNA and mRNA in the f1 to f6 subpopulations.** (A and B) HeLa cells were infected for 24 h with rICP47/vUs11 at an MOI of 5 and sorted into six subpopulations (f1 to f6) by cell sorting. Relative amounts of HSV-1 DNA (A) and mRNA (B) of selected genes in each subpopulation were analyzed by quantitative PCR or RNA-seq, respectively. Scatter plot of $\log_{10}$(relative MFI of Venus) vs $\log_{10}$(relative amount of HSV-1 DNA) (A), or scatter plots of $\log_{10}$(relative MFI of Venus) vs $\log_{10}$(relative FPKM) of the indicated HSV-1 genes (B) are shown. Each value is the mean ± standard error of the results of three biologically independent samples.

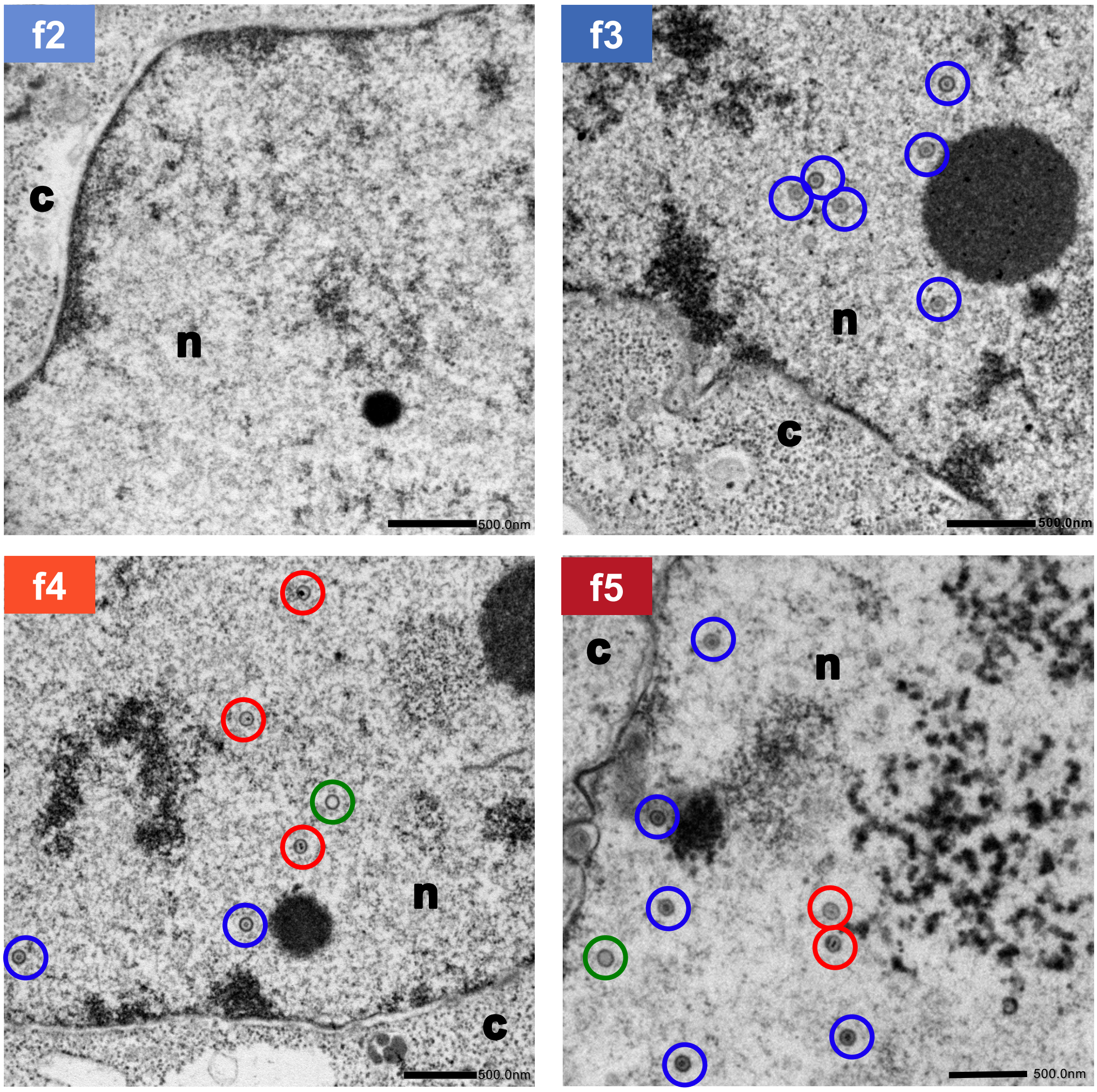


**Supplementary Fig. 8. Frequency of A, B, and C capsids in the nucleus of cells in f2 to f5 subpopulations.** A transmission electron microscopy image of cells in the f2 to f5 subpopulations. n, nucleus; c, cytoplasm from the experiment in Fig. 6. Type A capsids are marked in green, B capsids in blue, and C capsids in red. Scale bars = 500 nm.

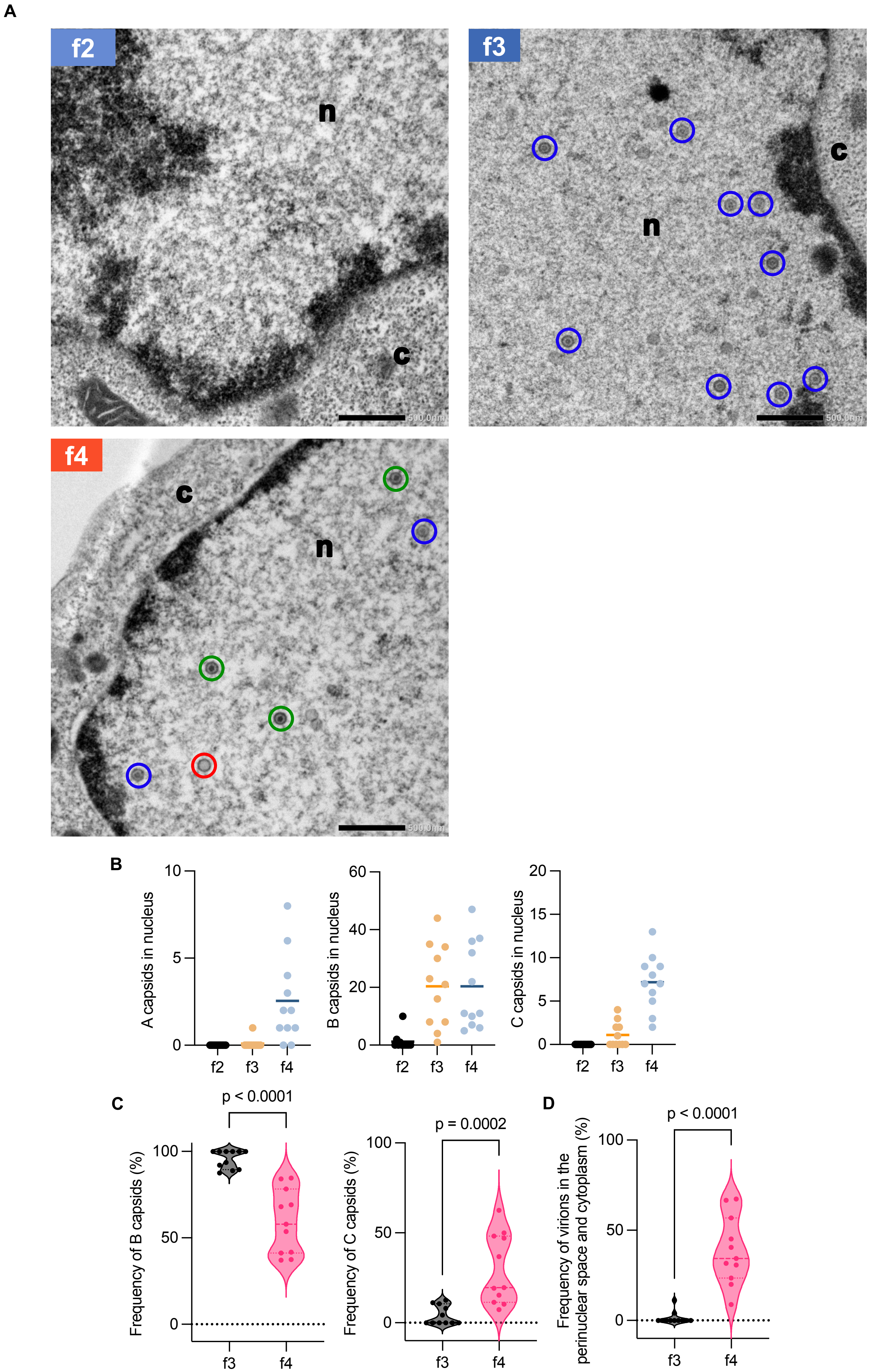


*(Legend on next page)*

**Supplementary Fig. 9 M. Nobe and Y. Maruzuru et al.**

**Supplementary Fig. 9. Frequencies of B and C capsids in the nucleus of cells in the f2 to f4 subpopulations at 8 h after infection.** (A) A transmission electron microscopy image of cells in the f2 to f4 subpopulations. n, nucleus; c, cytoplasm from the experiment in S-Fig. 6. Type A capsids are marked in green, B capsids in blue, and C capsids in red. Scale bar = 500 nm. (B) The numbers of A, B, and C capsids in the nucleus of 11 cells analyzed in S-Fig. 6 in the f2 to f4 subpopulations were quantitated. The horizontal bars indicate the means. Proportions of B and C capsids to nuclear capsids (C), or virions in the perinuclear space and cytoplasm (D) of cells with more than two nuclear capsids (f3, n = 11; f4, n = 11). Data are presented as the median ± interquartile range (IQR). Statistical analysis was performed by the Mann-Whitney *U*-test.

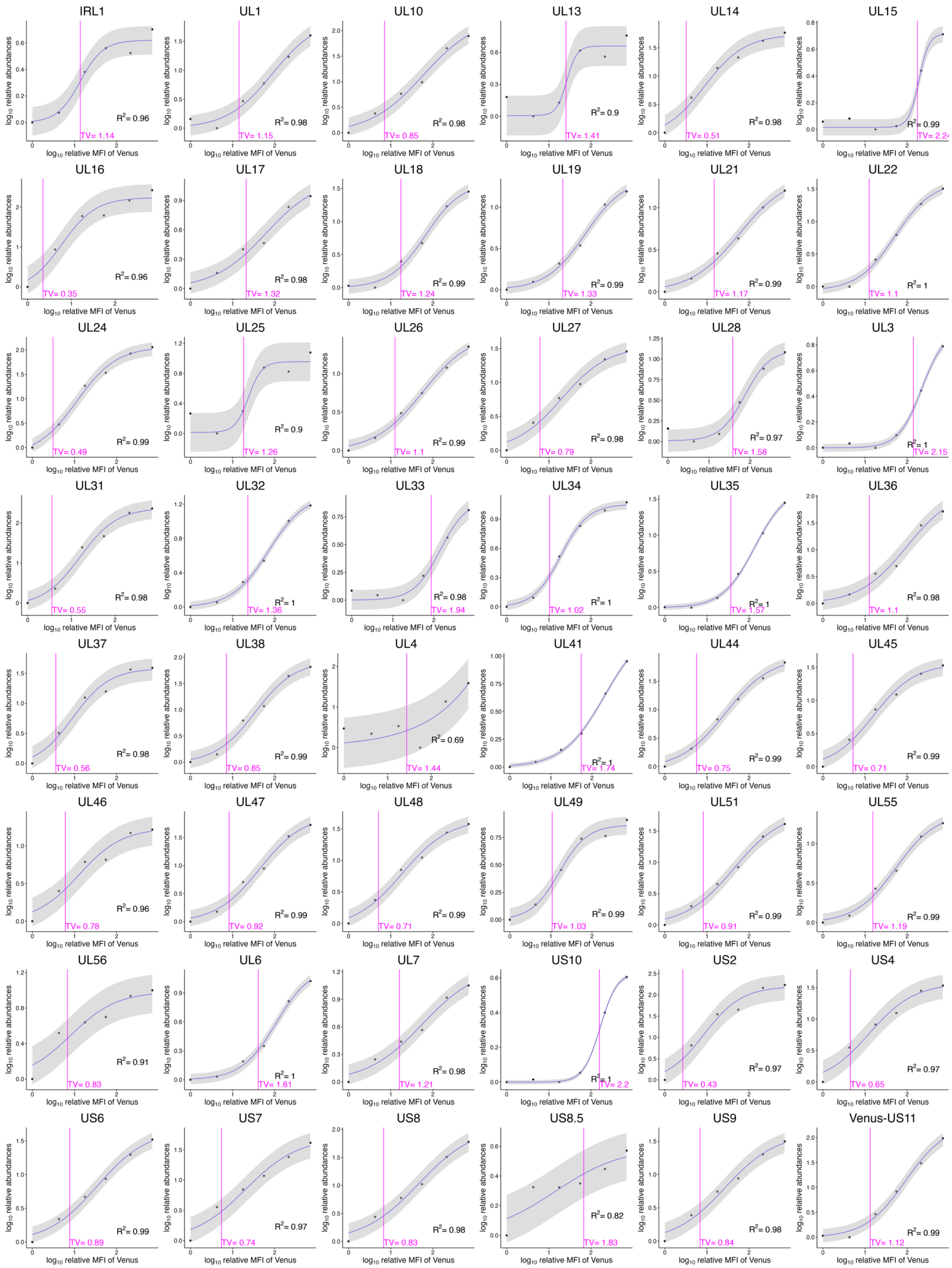


**Supplementary Fig. 10. Curve fitting to abundance profiles of HSV-1 L proteins.** Scatter plots of $\log_{10}$(relative MFI of Venus) vs $\log_{10}$(relative abundances) of the 48 indicated HSV-1 L proteins (from S-Fig. 1B) were fitted to a four-parameter logistic regression curve. TVs are the x-axis value at which the y-axis value increases by $\log_{10}2$ (2-fold increase in linear scale) from the starting point (x = 0) for each curve. The grey regions correspond to 95% confidence intervals. The average value of two biologically independent experiments was used for curve fitting.

**Supplementary Table 1. Oligonucleotide sequences, plasmid, and E. coli GS1873 containing HSV-BAC for the construction of recombinant viruses or mutagenesis.**

| Recombinant virus | Mutation inserted into pYEbac102Cre | Oligonucleotide sequence (5′-3′) | Plasmid DNA template | *E. coli* GS1873 containing HSV-BAC |
|---|---|---|---|---|
| YK410 | TagRFP-ICP47/ Venus-Us11 | 5'-AGTTTCTGGGCGGAGGGTGGTTCCCCCCGTGGCTCTCGAGATGGTGAGCAAGGGCGAGGA-3' | pBS-Venus-KanS (20) | *E. coli* GS1783/pYEbac102Cre |
| | | 5'-CGCCCGGCCCAACTGGGGCCGGGGGTTGGGTCTGGCTCATCTTGTACAGCTCGTCCATGC-3' | | |
| | | 5'-CCGTTGCGTGGACCGCTTCCTGCTCGTCGGGCGGGGGAGCATGGTGTCTAAGGGCGAAGA-3' | pFlag-TagRFP-KanS (This study) | *E. coli* GS1783/pYEbac102Cre carrying Venus fused to N-terminus of Us11 |
| | | 5'-TGTTGTCCAGGAAGGTGTCCGCCATTTCCAGGGCCCACGAATTAAGTTTGTGCCCCAGTT-3' | | |

**Supplementary Table 2. Number of cells and PFU attached to each subpopulation in Fig. 3D and E**

| Times after infection (h) | Subpopulation | Cell number of each subpopulation/$10^4$ cells of entire population | % of cells in each subpopulation | PFU in each subpopulation/$10^4$ cells of entire population | % of virus titer in each subpopulation |
|---|---|---|---|---|---|
| 4 | f1 | 9413.3 ± 216.3 | 94.1 ± 2.2 | 109.4 ± 38.8 | 95.9 ± 1.6 |
| 4 | f2 | 217.7 ± 110.4 | 2.2 ± 1.1 | 3.1 ± 0.4 | 4.1 ± 1.6 |
| 4 | f3 | N/A | N/A | N/A | N/A |
| 4 | f4 | N/A | N/A | N/A | N/A |
| 4 | f5 | N/A | N/A | N/A | N/A |
| 4 | f6 | N/A | N/A | N/A | N/A |
| 6 | f1 | 7871.7 ± 321.8 | 78.7 ± 3.2 | 32.2 ± 4.3 | 71.1 ± 6.7 |
| 6 | f2 | 1529.7 ± 261.5 | 15.3 ± 2.6 | 9.9 ± 2.3 | 21.7 ± 4.3 |
| 6 | f3 | 241.0 ± 123.6 | 2.4 ± 1.2 | 2.9 ± 1.3 | 7.1 ± 3.6 |
| 6 | f4 | N/A | N/A | N/A | N/A |
| 6 | f5 | N/A | N/A | N/A | N/A |
| 6 | f6 | N/A | N/A | N/A | N/A |
| 8 | f1 | 5427.3 ± 417.2 | 54.3 ± 4.2 | 13.2 ± 2.1 | 6.1 ± 1.3 |
| 8 | f2 | 2405.0 ± 145.0 | 24.1 ± 1.4 | 13.8 ± 1.4 | 7.8 ± 2.7 |
| 8 | f3 | 1367.3 ± 132.8 | 13.7 ± 1.3 | 37.9 ± 0.2 | 20.3 ± 5.8 |
| 8 | f4 | 410.3 ± 132.7 | 4.1 ± 1.3 | 162.3 ± 69.9 | 55.0 ± 3.0 |
| 8 | f5 | N/A | N/A | N/A | N/A |
| 8 | f6 | N/A | N/A | N/A | N/A |
| 10 | f1 | 3458.3 ± 308.2 | 34.6 ± 3.1 | 27.8 ± 5.9 | 2.4 ± 1.0 |
| 10 | f2 | 2114.3 ± 63.9 | 21.1 ± 0.6 | 18.3 ± 1.9 | 1.3 ± 0.3 |
| 10 | f3 | 2259.7 ± 82.5 | 22.6 ± 0.8 | 96.3 ± 15.9 | 6.7 ± 1.5 |
| 10 | f4 | 1245.0 ± 108.7 | 12.5 ± 1.1 | 430.3 ± 64.7 | 34.3 ± 10.4 |
| 10 | f5 | 559.0 ± 115.4 | 5.6 ± 1.2 | 1080.4 ± 518.6 | 52.7 ± 10.6 |
| 10 | f6 | N/A | N/A | N/A | N/A |
| 12 | f1 | 2528.3 ± 342.5 | 25.3 ± 3.4 | 31.6 ± 4.1 | 1.0 ± 0.4 |
| 12 | f2 | 1564.3 ± 74.4 | 15.6 ± 0.7 | 38.9 ± 14.1 | 0.7 ± 0.2 |
| 12 | f3 | 2193.7 ± 91.3 | 21.9 ± 0.9 | 108.7 ± 16.6 | 3.4 ± 1.7 |
| 12 | f4 | 1725.0 ± 142.1 | 17.3 ± 1.4 | 857.3 ± 100.7 | 20.4 ± 4.2 |
| 12 | f5 | 1392.0 ± 262.8 | 13.9 ± 2.6 | 3279.4 ± 942.7 | 62.2 ± 5.3 |
| 12 | f6 | 209.3 ± 54.7 | 2.1 ± 0.5 | 649.8 ± 197 | 12.3 ± 1.2 |
| 24 | f1 | 1215.0 ± 161.8 | 12.2 ± 1.6 | 105.0 ± 7.2 | 1.5 ± 0.1 |
| 24 | f2 | 889.3 ± 63.2 | 8.9 ± 0.6 | 74.0 ± 9.6 | 1.1 ± 0.1 |
| 24 | f3 | 2151.0 ± 121.7 | 21.5 ± 1.2 | 145.2 ± 25.8 | 2.1 ± 0.3 |
| 24 | f4 | 1939.0 ± 139.1 | 19.4 ± 1.4 | 580.5 ± 135.4 | 8.3 ± 1.5 |
| 24 | f5 | 1695.3 ± 58.7 | 17.0 ± 0.6 | 2282.8 ± 267.6 | 33.8 ± 4.0 |
| 24 | f6 | 1680.3 ± 145.7 | 16.8 ± 1.5 | 3728.9 ± 707.1 | 53.1 ± 5.7 |

Values are the mean ± SE. N/A, not available.
